# Enhanced endogenous activity predicts elimination of adult-born neurons in the mouse olfactory bulb

**DOI:** 10.1101/2020.09.09.286591

**Authors:** Xin Su, Yury Kovalchuk, Nima Mojtahedi, Olga Garaschuk

## Abstract

Adult-born cells, arriving daily into the rodent olfactory bulb, either integrate into the neural circuitry or get eliminated. Whether these two populations differ in their morphological or functional properties remains, however, unclear. Using longitudinal *in vivo* two-photon imaging, we monitored dendritic morphogenesis, odor-evoked responsiveness, endogenous Ca^2+^ signaling, and survival/death of adult-born juxtaglomerular neurons (abJGNs). We found that maturation of abJGNs is accompanied by a significant reduction in dendritic complexity, with surviving and subsequently eliminated cells showing similar degrees of dendritic remodeling. Surprisingly, 63% of eliminated abJGNs acquired odor-responsiveness before death, with amplitudes and time courses of odor-evoked responses similar to those recorded in surviving cells. We observed, however, a significant long-lasting enhancement of endogenous Ca^2+^ signaling in subsequently eliminated abJGNs, visible already 6 days before death. These findings identify ongoing endogenous Ca^2+^ signaling as a key predictor of abJGNs’ fate.

## Introduction

Adult neurogenesis provides the brains of grown-ups with an additional level of plasticity and cognitive flexibility (Doetsch and Hen, 2005; Ming and Song, 2011), which is important for odor-dependent learning, memory encoding, spatial, temporal or odor pattern separation and emotional control (Alonso et al., 2012; Anacker and Hen, 2017). Altered adult neurogenesis has been associated with several neurodegenerative (e.g. Parkinson’s, Alzheimer’s and Huntington’s disease) and neuropsychiatric disorders, including major depressive disorder, epilepsy, schizophrenia and anxiety (Braun and Jessberger, 2014; Gonçalves et al., 2016b; Parent and Murphy, 2008).

Under physiological conditions, two regions of the adult rodent brain continuously generate new neurons: the subgranular zone of the dentate gyrus, generating hippocampal granule cells, and the subventricular zone (SVZ) of the lateral ventricle, generating adult-born cells (ABCs) of the olfactory bulb (OB) (Denoth-Lippuner and Jessberger, 2021; Ming and Song, 2011). Neuronal progenitors generated in the SVZ migrate through the rostral migratory stream (RMS) into the OB, where they differentiate into local inhibitory interneurons. Approximately 90% of ABCs become granule cells (GCs), whereas another 5-10% become juxtaglomerular neurons (JGNs; Alvarez-Buylla and García-Verdugo, 2002; Bonfanti and Peretto, 2011). It takes approximately 4-6 days for ABCs to migrate from the SVZ into the core of the OB, where they start a saltatory (i.e. ‘stop-and-go’-like) radial migration (Liang et al., 2016). Whereas GCs stop at the end of the radial migration phase, abJGNs speed up and, after arrival into the glomerular layer, switch from the radial to the long-distance, uni- or multi-directional lateral migration, lasting till the end of the pre-integration phase (∼3-4 weeks after ABCs’ birth) (Li et al., 2021; Liang et al., 2016).

The patterns of morphogenesis also differ between adult-born GCs and JGNs. Whereas adult-born GCs start to elaborate their dendritic trees after arrival to the final destination (Sailor et al., 2016), abJGNs migrate within the glomerular layer with elaborated albeit constantly changing dendritic trees (Kovalchuk et al., 2015). Longitudinal *in vivo* imaging showed that ∼30 days after birth, adult-born GCs gain stable dendritic morphology (Sailor et al., 2016). For abJGNs, so far the development of their dendritic tree was only studied at the population level. The cells increased their complexity between 10 and 45 days after birth, reaching the steady-state thereafter (Livneh et al., 2009; Mizrahi, 2007). Still, even after maturation, the structural dynamics of dendrites and spines of both adult-born GCs and JGNs remains high, suggesting that ABCs are the most plastic neuronal population in the OB (Mizrahi, 2007; Sailor et al., 2016).

Thousands of ABCs migrate into the OB every day but many of them are thought to be eliminated by programmed cell death (Biebl et al., 2005, 2000; Petreanu and Alvarez-Buylla, 2002; Whitman and Greer, 2007; Winner et al., 2002; Woon et al., 2007), but see (Platel et al., 2019). According to data obtained in bromodeoxyuridine (BrdU) incorporation experiments, elimination of ABCs mainly happens between 15 and 45 days after birth (Petreanu and Alvarez-Buylla, 2002; Whitman and Greer, 2007; Winner et al., 2002). Plenty of factors, including intracellular signaling molecules and pathways (pro- and anti-apoptotic Bcl-2 family, cAMP response element-binding protein signaling pathway), extracellular signaling molecules (e.g. neurotrophic and growth factors, hormones) and neurotransmitters (GABA, glutamate, serotonin, acetylcholine) were shown to influence the survival of ABCs (Benn and Woolf, 2004; Fomin-Thunemann and Garaschuk, 2021; Khodosevich et al., 2013; Kuhn, 2015). The latter is also modulated by cellular components of the brain’s immune system, including microglia and astrocytes (Sierra et al., 2010; Sultan et al., 2015).

The survival of ABCs appears to depend on odor sensation, as enriched odor exposure and olfactory discrimination learning promote while odor deprivation (by closing one nostril or ablating olfactory sensory neurons) inhibits the survival of ABCs (Alonso et al., 2006; Corotto et al., 1994; Sawada et al., 2011; Sultan et al., 2011). Previous reports suggest the existence of a critical period (∼2-4 weeks after birth) when ABCs are most sensitive to the influence of sensory experience (Alonso et al., 2006; Mouret et al., 2008; Yamaguchi and Mori, 2005). Since this period corresponds to the pre-integration phase during which ABCs accomplish their migration and start receiving synaptic inputs, it has been postulated that ABCs are eliminated because they “fail to integrate” into the pre-existing neuronal circuitry (Lin et al., 2010; Turnley et al., 2014). However, this hypothesis has never been tested directly. It remains also unclear whether there are any fundamental morphological or functional differences between ABCs that survive and those ABCs that become eliminated. In this study, we employed longitudinal *in vivo* single-cell tracking and ratiometric two-photon Ca^2+^ imaging to test whether the surviving and the subsequently eliminated abJGNs differ in terms of dendritic morphology, and endogenous or sensory-driven Ca^2+^ signaling. We also examined whether abJGNs are eliminated because they fail to integrate into the pre-existing neural circuitry.

## Results

### Longitudinal *in vivo* monitoring of fate selection of adult-born juxtaglomerular neurons

For unequivocal identification of migrating abJGNs in longitudinal experiments, we used the red-green-blue (RGB) cell-marking approach (Fig. 1a), labeling ABCs via an RMS injection of a mixture of retroviruses encoding mCherry (red), Venus (green) and Cerulean (blue) (Liang et al., 2016). This technique provides each labeled cell with a defined color identity, thus enabling accurate tracking of migrating cells over the entire experiment (Gomez-Nicola et al., 2014; Liang et al., 2016). The RGB-labeled abJGNs were visualized through a cranial window in awake mice and their position as well as dendritic morphology were monitored longitudinally starting at 12 days post-viral injection (DPI 12) till DPI 45 (Fig. 1b,c). To develop criteria for the detection of cell elimination, we made use of our prior results showing that cells rarely moved after being stable for more than 4 days, see Figure 4 in (Liang et al., 2016), and left a 30-50-µm-wide margin to the border of the field of view (FOV), not to mistake cell migration out of the FOV for cell death. Thus, abJGNs were scored as eliminated if (i) cell debris was found in the position where an adult-born JGN was previously located (upper panel in Fig. 1d) or (ii) a cell was present within the FOV at DPI 12, stayed in the same position at DPI 18-22 (at least 4 days) and then disappeared from the dorsal surface of the bulb under the entire cranial window (lower panel in Fig. 1d). At DPI 12-22, cells were scored eliminated only if their debris was found. We often noticed that debris of cells expressing mCherry together with other fluorescent proteins was colored red (e. g. magenta arrow in Fig. 1d, upper panel), likely due to the higher resistance of mCherry to proteolysis and degradation (Costantini et al., 2015). Based on the above criteria, RGB-labeled abJGNs were classified as surviving (i.e. staying in the same FOV from DPI 12 to DPI 45), eliminated or uncertain (disappearing from the FOV without fulfilling the above criteria) cells. To avoid false-positive results, we excluded the uncertain cells from further analyses. Note that this might cause an underestimation of the fraction of eliminated cells, reported in this study.

**Figure 1.**
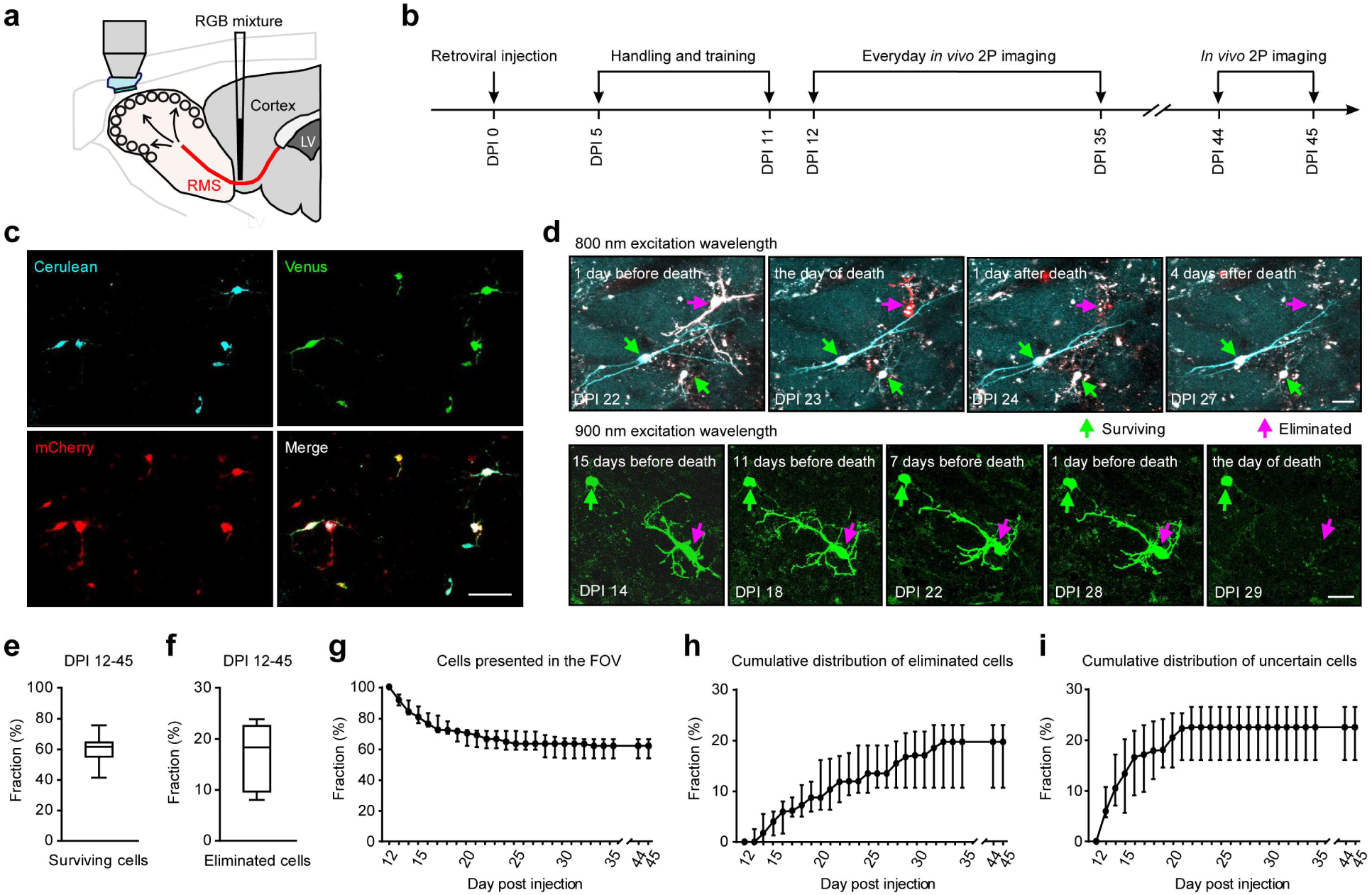
Longitudinal tracking of abJGNs’ fate. (**a**) Scheme of the experimental setup. AbJGNs were labeled via RMS injection of a 1:1:1 mixture of RGB retroviruses encoding Venus (green), mCherry (red) and Cerulean (blue) fluorescent proteins and repeatedly imaged via a chronic cranial window using two-photon (2P) microscopy. (**b**) Illustration of the experimental timeline. (**c**) Maximum intensity projection (MIP) images (0-100 μm, step 2 μm) of RGB^+^ abJGNs taken at DPI 12 in red, green and blue channels. The merged image is shown in the lower right corner. Scale bar: 50 µm. (**d**) Upper: MIP images (34-54 µm, step 2 µm, 800 nm excitation wavelength) showing 1 eliminated cell (magenta arrow) and 2 surviving cells (green arrows) in the same FOV at 4 different time points: DPI 22 (1 day before death), DPI 23 (the day of death), DPI 24 (1 day after death) and DPI 27 (4 days after death). Note red cell debris visible at DPI 23 and DPI 24. Lower: MIP images (12-52 µm, step 2 µm, 900 nm excitation wavelength) illustrating 1 eliminated and 1 surviving abJGNs in the same FOV at 5 different time points. The day of death was DPI 29. Scale bars: 25 µm. (**e, f**) Box plots showing the median (per mouse) fractions of surviving (**e**) and eliminated (**f**) abJGNs at DPI 12-45 (n = 12 mice). (**g**) Graph showing the fraction of RGB^+^ abJGNs present within FOVs at different DPIs. The number of cells present at DPI 12 is taken as 100%. (**h,i**) Graphs showing cumulative fractions of eliminated (**h**) and uncertain (**i**) cells at DPI 12-45 (n = 12 mice).

With the above criteria imposed, 61.90 ± 11.29% (per mouse) of 528 analyzed abJGNs were classified as surviving cells (Fig. 1e, n = 12 mice), 18.45 ± 12.89% as eliminated cells (Fig. 1f) and the rest as uncertain cells. The cell turnover within the FOVs was fast at DPI 12-25 and substantially slower at later time points (Fig. 1g), with the cumulative fraction of eliminated cells increasing almost linearly from DPI 12 to DPI 34 (Fig. 1h; 91 cells, n = 12 mice) and the cumulative fraction of uncertain cells (Fig. 1i) mirroring that of migrating cells in our previous study, see Figure 7 in (Liang et al., 2016). In total, 91.21% of eliminated abJGNs (83/91 cells) died between DPI 12 and DPI 28, with no cells being eliminated after DPI 34. These data are consistent with a previous study showing that the elimination of abJGNs after the end of the pre-integration phase (day 28) becomes minimal (Sawada et al., 2011).

Taken together, our intravital imaging data clearly documented the death of at least ∼20% of abJGNs. The death was cell age-dependent, mainly happening within the first 34 days after cells’ birth.

### Longitudinal analyses of dendritic development of adult-born JGNs

We next aimed to identify features that may predict the fate of abJGNs. At the population level, previous work showed that dendritic trees of abJGNs are more complex at DPI 43-45 compared to DPI 10-13 (Livneh et al., 2014, 2009; Mizrahi, 2007), but longitudinal data from individual cells are lacking. Therefore, we followed individual cells over time and analyzed their dendritic morphology, based on total dendritic branch length (TDBL), number of dendrites, number of primary dendrites, number of branch points, and number of dendritic endings (Fig. 2a). Based on the number of primary dendrites, abJGNs were classified as either unipolar or multipolar cells (Fig. 2b). We found a significant increase in the fraction of unipolar abJGNs at DPI 45, compared to DPI 13 (Fig. 2c, *P* = 4.9*10^-3^, Wilcoxon signed-rank test, n = 11 mice). There are two possible explanations for this result: (i) either multipolar cells pruned their primary dendrites to become unipolar cells or (ii) the unipolarity of the dendritic tree is beneficial for cell survival and more multipolar cells were eliminated at DPI 13-45. Out of all abJGNs with documented dendritic morphology, 22.5% (9/40 cells, n = 11 mice) of eliminated and 12.16% of surviving cells (18/148 cells, n = 12 mice) had a unipolar morphology at DPI 13. The observed difference, however, did not reach the level of statistical significance (Fig. 2d, *P* = 0.16, Chi-square test with Yates’ correction), suggesting that the unipolarity of the dendritic tree is not beneficial for the survival of abJGNs.

**Figure 2.**
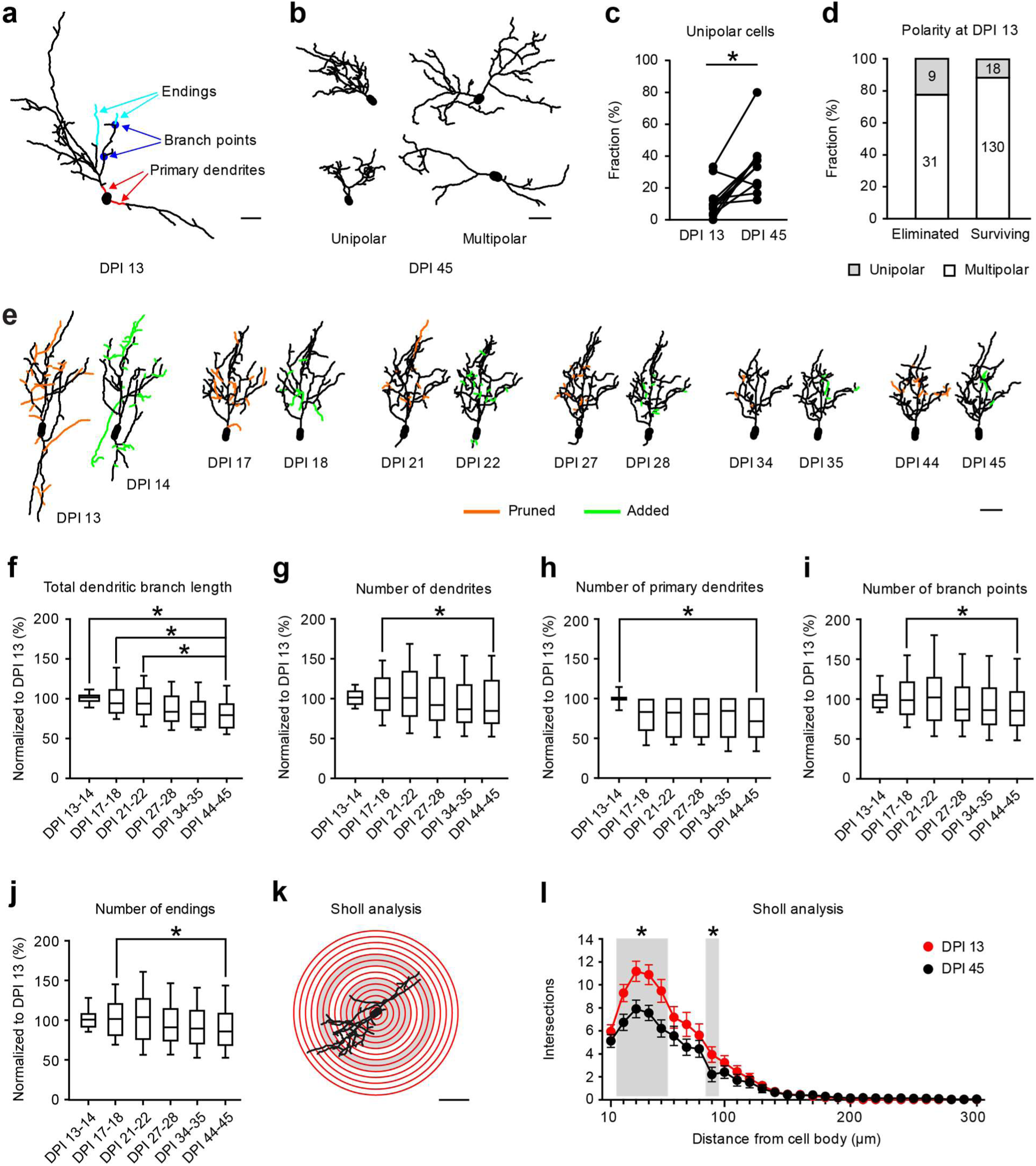
Dendritic development of abJGNs. (**a**) 2D projection image of a reconstructed adult-born JGN (DPI 13), exemplifying the morphological parameters analyzed in this study. Primary dendrites are color-coded red, some branch points blue and some dendritic endings cyan. Scale bar: 25 µm. (**b**) Representative reconstructions of 2 unipolar and 2 multipolar cells at DPI 45. Scale bar: 25 µm. (**c**) Connected dot graph showing the fractions of unipolar cells at DPI 13 and DPI 45 (P = 4.9*10^-3^, Wilcoxon signed rank test; n = 11 mice). (d) Bar graph showing the fractions of unipolar and multipolar cells among the eliminated (left) and surviving (right) cells (*P* = 0.16, Chi-square test with Yates’ correction). Cell polarity was evaluated at DPI 13. (**e**) Representative reconstructions of the same cell at 12 different time points (groups of 2 consecutive time points, DPI 13-45). Colors highlight pruned (orange) and added (green) dendrites for each group. Scale bar: 25 µm. (**f-j**) Box plots illustrating the normalized total dendritic branch length (**f**) and the numbers of dendrites (**g**), primary dendrites (**h**), branch points (**i**) and endings (**j**). For each cell, the data were normalized to the respective value measured at DPI 13 (*P* < 0.05, Friedman test followed by Dunn’s multiple comparisons test; n = 34 cells, 8 mice). (**k**) Schematic diagram illustrating the Sholl analysis procedure. Scale bar: 30 μm. (**l**) Graph comparing the mean numbers of intersections with Sholl spheres for cells imaged at DPI 13 (red) and DPI 45 (black). Error bars: SEM. Grey zones in (**k**) and (**l**) indicate the regions where the numbers of intersections showed significant differences between the two age groups (\**P* < 0.05, Wilcoxon signed rank test; n = 34 cells, 8 mice). The following supplementary figures are related to Figure 2: **Supplementary Figure 1**. 4D structural plasticity analysis of dendritic remodeling in stable surviving abJGNs. **Supplementary Figure 2**. Development and plasticity of the dendritic tree in stable surviving adult-born PGCs. **Supplementary Figure 3**. Development and plasticity of the dendritic tree in stable surviving adult-born SACs. **Supplementary Figure 4**. Initial period of rapid dendritic growth in surviving abJGNs at DPI 9-13. **Supplementary Figure 5**. Impact of the improved image quality on the dendritic complexity of abJGNs.

Next, we analyzed the dendritic morphology of surviving abJGNs, which did not change their position from DPI 13 to DPI 45 (called stable surviving abJGNs; 69.62 ± 12.12% of all surviving cells, n = 12 mice). We reconstructed and analyzed the dendritic trees of stable surviving abJGNs at 12 different time points as indicated in Fig. 2e. For each cell, the data were normalized to the respective value measured at DPI 13 and means of 2 neighboring time points were used for statistics shown in Fig. 2f-j. Unexpectedly, all morphological parameters of stable surviving abJGNs including the median (per mouse) TDBL, number of dendrites, number of primary dendrites, number of branch points and number of endings declined with cells’ maturation (Fig. 2f-j, n = 34 cells, 8 mice). The reduction in the number of primary dendrites is consistent with the increased fraction of unipolar cells seen at DPI 45 (Fig. 2c). Concordantly, Sholl analysis also revealed a significant reduction of the dendritic arbor complexity at DPI 45 compared to DPI 13 (Fig. 2k,l).

To investigate the turnover of dendritic branches, we conducted quantitative 4-dimensional structural plasticity analyses (4DSPA), using published protocols (Gonçalves et al., 2016a; Lee et al., 2013). For each neuron, the degree of dendritic remodeling (i.e. the fractions of pruned or added dendritic endings) was analyzed at 6 different time points (DPI 13-14, DPI 17-18, DPI 21-22, DPI 27-28, DPI 34-35 and DPI 44-45; Fig. 2e). The data showed that stable surviving abJGNs underwent extensive dendritic remodeling at early time points and the degree of remodeling decreased over time (Supplementary Fig. 1). Consistent with the literature data (Livneh and Mizrahi, 2011; Mizrahi, 2007), even at DPI 44-45, abJGNs remained structurally dynamic (Fig. 2e, Supplementary Fig. 1).

Based on the pattern of the expressed molecular markers, morphological, electrophysiological and functional properties, abJGNs are further subdivided into two distinct subtypes: periglomerular cells (PGCs) and short axon cells (SACs) (Kosaka and Kosaka, 2011; Nagayama et al., 2014). PGCs arborize in a single glomerulus and have short but complex branches, while SACs arborize in several glomeruli and branch less frequently (Bywalez et al., 2017; Kiyokage et al., 2010; Nagayama et al., 2014; Pinching and Powell, 1971). Based on their dendritic tree at DPI 45, we classified the abJGNs into PGCs (43.75 ± 10.00%), SACs (46.67 ± 20.83%) and cells with unclear morphology (6.25 ± 18.75%; n = 11 mice). For stable surviving PGCs (Supplementary Fig. 2, n = 13 PGCs, 7 mice) and SACs (Supplementary Fig. 3, n = 14 SACs, 5 mice), the major aspects of their morphological development and dendritic remodeling were similar. Both cell types exhibited a significant reduction in the TDBL as well as the number of primary dendrites from DPI 13 to DPI 45. For both subtypes, we also observed a decrease in the degree of dendritic remodeling during maturation. Thus, despite distinct dendritic morphology, adult-born PGCs and SACs share similar patterns of dendritic development.

In a subset of experiments, we additionally followed the dendritic morphology of abJGNs from DPI 9 to DPI 13. The dendritic morphology of stable surviving abJGNs (i.e. cells, not changing their position at DPI 13-45) was analyzed regardless of whether they migrated or not at DPI 9-13. All the data were normalized to the respective values measured at DPI 13. These data revealed that dendritic trees of stable surviving abJGNs grew rapidly at DPI 9-13, increasing, for example, the TDBL by 35.21% and the number of dendrites by 68.42% between DPI 9 and DPI 13 (Supplementary Fig. 4, n = 15 cells, 4 mice). To test whether settings optimized for the longitudinal imaging (i.e. high photomultiplier gain, low laser power; Supplementary Fig. 5a) lead to underestimation of dendritic complexity, in a control experiment we optimized settings for high image quality (i.e. lower photomultiplier gain, higher laser power; Supplementary Fig. 5b) and directly compared the two sets of data. In the low-noise group, the signal-to-noise ratio (SNR) increased ∼3 times (291.32 ± 137.63%, *P* < 0.001, n = 12 cells; here and below Wilcoxon signed-rank test), but the signal-to-background ratio (SBR) did not change significantly (98.47 ± 39.95%, *P* = 0.79; Supplementary Fig. 5c). In general, the improvement of image quality did not change the dendritic complexity of abJGNs (Supplementary Fig. 5d). The number of dendritic endings, however, was slightly but significantly higher (105.20 ± 6.23%, P < 0.001).

Taken together, our data revealed that in stable surviving abJGNs, dendritic growth was restricted to a short initial time window (DPI 9-13). The rest of the maturation time (DPI 13-45) was accompanied by a significant reduction in the size and complexity of the dendritic tree. At early developmental stages, abJGNs underwent both extensive dendritic pruning and addition. The degree of dendritic remodeling decreased during development, with adult-born PGCs and adult-born SACs sharing similar developmental patterns.

### Similar levels of dendritic complexity and morphological plasticity in eliminated and surviving abJGNs

Elimination of immature neurons by apoptosis, taking place both during development and in adulthood (Biebl et al., 2005; Kuhn, 2015; Yuan and Yankner, 2000), is a fast process, as the time from the initiation of apoptosis to its completion usually takes less than 24 hours (Cellerino et al., 2000; Cordeiro et al., 2004; Elmore, 2007; Saraste, 1999). To determine whether the morphological features can distinguish subsequently eliminated from surviving cells, we took the day of cell death as a reference point and analyzed the morphology of eliminated abJGNs at one, two and three days before death (DBD 1: 0-24 hours before death, DBD 2: 24-48 hours before death, DBD 3: 48-72 hours before death). The addition and pruning of dendritic endings between two neighboring time points were investigated using the 4DSPA approach. The data obtained were normalized to the respective median values of all (at least 4) surviving cells of the same subtype (i.e. either PGCs-like or SACs-like) recorded in the same mouse at the same time point, as illustrated in Fig. 3a. The terms PGCs- or SACs-like reflect the fact that the cells’ dendritic morphology can be unequivocally determined only in the mature state. Interestingly, even at 1-2 days before death, the subsequently eliminated cells continued to prune and add dendritic endings and we found no significant difference between eliminated and surviving abJGNs for all the morphological parameters analyzed, including TDBL (Fig. 3b) and the number of dendrites (Fig. 3c), number of primary dendrites (Fig. 3d), number of branch points (Fig. 3e), number of endings (Fig. 3f) as well as the fraction of pruned (Fig. 3g) and added (Fig. 3h) endings (*P* > 0.05 for all comparisons, Wilcoxon signed rank test; n = 10 eliminated and 63 surviving cells, 6 mice). Thus, our data showed that the complexity of the dendritic tree and the remodeling dynamics cannot predict the subsequent elimination of abJGNs.

**Figure 3.**
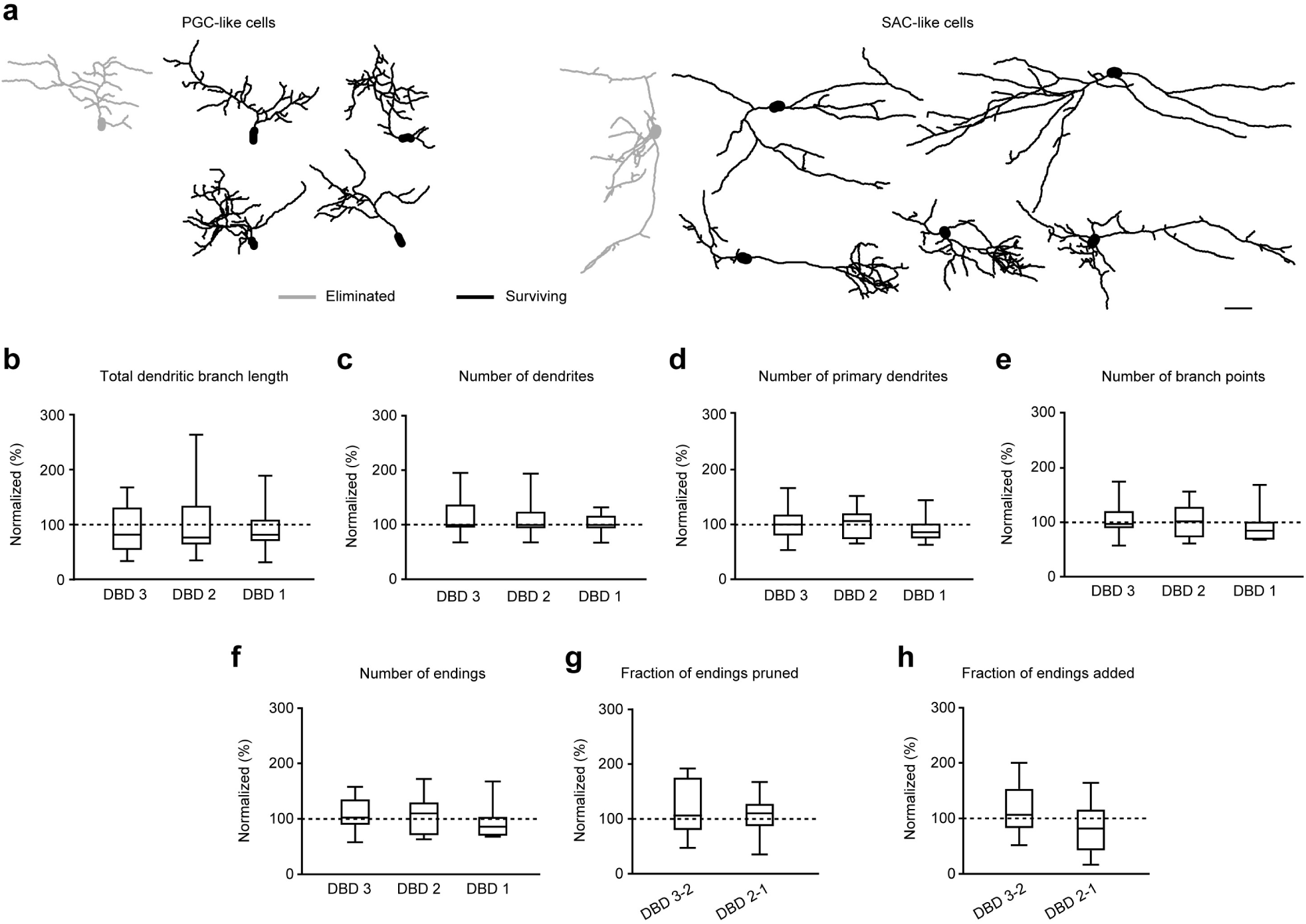
Dendritic morphology cannot predict the fate of abJGNs. (**a**) Representative reconstructions of eliminated (grey) and corresponding surviving (black) abJGNs from the same mice at DBD 2. Left: PGC-like cells, right: SAC-like cells. Scale bar: 25 µm. (**b-f**) Box plots showing the normalized total dendritic branch length (**b**), numbers of dendrites (**c**), primary dendrites (**d**), branch points (**e**) and endings (**f**) measured at DBD 3, DBD 2 and DBD 1. (**g, h**) Normalized fractions of pruned (**g**) and added (**h**) dendritic endings at DBD 3-2 and DBD 2-1. Data obtained in subsequently eliminated cells were normalized to the respective median value obtained in corresponding surviving cells of the same type recorded in the same animal at the same time point (*P* > 0.05, Wilcoxon signed rank test; n = 10 eliminated cells and 63 surviving cells, 6 mice).

**Figure 4.**
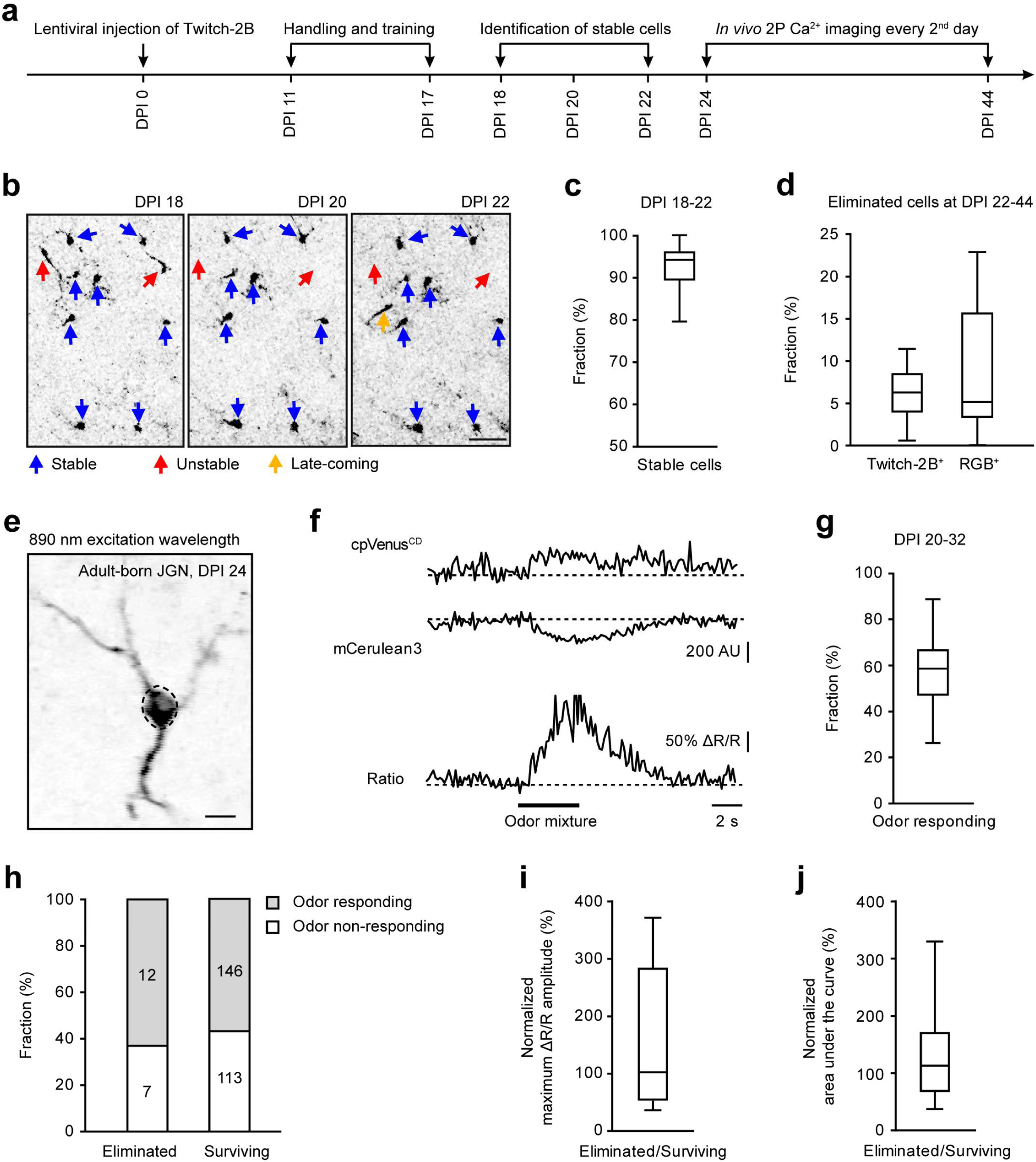
Fate of abJGNs is not determined by their odor-responsiveness. (**a**) Schematic diagram of experimental timeline. (**b**) MIP (63-135 µm, step 3 µm) images showing stable (blue arrows), unstable (red arrows) and late-coming (orange arrow) Twitch-2B^+^ abJGNs at DPI 18-22. Scale bar: 50 µm. (**c**) Box plot illustrating the fraction of stable cells at DPI 18-22 (n = 11 mice). (**d**) Box plot showing the fractions of cells eliminated at DPI 22-44 in Twitch-2B^+^ and RGB^+^ cell groups (*P* = 0.97, Mann-Whitney test; n = 11 and 12 mice, respectively). (**e**) Representative image (average of 140 consecutive frames) of a Twitch-2B^+^ abJGN at DPI 24. Scale bar: 10 μm. (**f**) Traces illustrating changes in fluorescence intensity of mCerulean3, cpVenus^CD^ and the Twitch-2B ratio trace recorded from the somatic region of interest shown in (**e**) in response to a 4-s-long application of the mixture of 7 odorants (ethyl-acetate, butanal, pentanal, ethyl tiglate, propanal, methyl-propionate and ethyl-butyrate). (**g**) Box plot summarizing the fraction of odor responding cells at DPI 20-32 (n = 11 mice). (**h**) Bar graphs showing the fractions of odor responding and odor non-responding cells among the eliminated (left) and surviving (right) cells (*P* = 0.74, Chi-square test with Yates’ correction). (**i,j**) Box plots showing the normalized maximum ΔR/R amplitude (**i**) (*P* = 0.49, Wilcoxon signed rank test) and area under the curve (**j**) (*P* = 0.23, One sample *t* test). The medians of subsequently eliminated cells were normalized to the medians of corresponding surviving cells recorded in the same animal at the same time point (n = 12 odor responding eliminated and 112 odor responding surviving cells, 8 mice). The following supplementary figure is related to Figure 4: **Supplementary Figure 6**. Visualization of the death of Twitch-2B^+^ abJGNs.

### Can odor-responsiveness predict abJGNs’ fate?

Since the critical period of ABCs’ elimination corresponds to the pre-integration phase, when ABCs start to receive synaptic inputs and integrate into the local circuitry, it has been postulated that ABCs die because they “fail to integrate” into the pre-existing neural circuitry (Lepousez and Lledo, 2011; Lin et al., 2010; Turnley et al., 2014). We took odor-responsiveness as the readout of adult-born JGNs’ functional integration into the neural circuitry and tested whether odor-responsiveness protects abJGNs from being eliminated. To this end, migrating ABCs in the RMS were labeled by lentivirus encoding the FRET-based Ca^2+^ indicator Twitch-2B (Thestrup et al., 2014). Because of its ratiometric nature, Twitch-2B is particularly valuable for longitudinal imaging of individual cells in awake animals (Maslyukov et al., 2018; Thestrup et al., 2014). Since it is difficult to determine the identity of migrating abJGNs without unique color tags, our longitudinal imaging started at DPI 18 (Fig. 4a), when the majority of abJGNs have completed their lateral migration (see Figure 7A in Liang et al., 2016). We focused on the cells, which did not change their position during the first three consecutive imaging sessions (DPI 18, 20 and 22), referred to as stable abJGNs (Fig. 4b). Focusing on stable, post-migratory abJGNs also helped to avoid late-coming cells (yellow arrow in Fig. 4b), arriving in the OB at a low rate after lentiviral injection in the RMS (Mizrahi, 2007; Wallace et al., 2017). Under these recording conditions, 94.12 ± 7.10% (median per mouse) of Twitch-2B^+^ abJGNs were stable cells (Fig. 4c, n = 533 cells, 11 mice). These cells were imaged further every 2^nd^ day in awake mice (Fig. 4a).

Similar to the criteria described for RGB-labeled cells (see above), Twitch-2B^+^ abJGNs were scored as eliminated either when the cell debris was identified or when stable cells disappeared after DPI 22 (Supplementary Fig. 6). The median (per mouse) fraction of Twitch-2B^+^ abJGNs eliminated from DPI 22 to DPI 44 was 6.25 ± 5.24% (n = 11 mice) and thus similar to the fraction (5.10 ± 9.87%, n = 12 mice) of RGB^+^ abJGNs eliminated during the same time period (Fig. 4d, *P* = 0.97, Mann-Whitney test). To test whether longitudinal imaging might enhance the elimination of abJGNs, we compared the fractions of surviving cells in our RGB and Twitch-2B group to a dataset from our laboratory (Li et al., 2021), in which Twitch-2B^+^ abJGNs were imaged only at 2 time points: either at DPI 14 and 25 or DPI 25 and 45. We did not detect any significant difference both at DPI 14/25 (2-time-points group: 62.50 ± 23.41%, n = 5 mice; RGB group: 61.18 ± 24.24%, n = 12 mice; *P* = 0.65, Mann-Whitney test) and DPI 25/45 (2-time-points group: 80.00 ± 20.76%, n = 5 mice; RGB group: 96.29 ± 7.33%, n = 12 mice; Twitch-2B group: 94.74 ± 4.48%, n = 11 mice; *P* = 0.07, Kruskal-Wallis test), suggesting that our longitudinal imaging protocol did not promote the elimination of abJGNs.

Odor-responsiveness of Twitch-2B^+^ abJGNs was examined by applying the mixture of either 3 or 7 odorants (see Methods) in front of the snout of anesthetized mice. Both odor mixtures cause broad activation of dorsal glomeruli in the OB (Kovalchuk et al., 2015; Livneh et al., 2014), thereby increasing the probability to recruit odor responding abJGNs. Under these experimental conditions, 58.00 ± 17.44% (median per mouse, n = 281 cells tested in 11 mice) of abJGNs showed odor-evoked responses (Fig. 4e-g). Out of 25 subsequently eliminated cells, the odor-evoked responsiveness of 19 cells happened to be tested before they were eliminated. Out of these 19 cells, 63.16% of cells (12/19 cells, 8 mice) showed odor-evoked Ca^2+^ signals prior to death (Fig. 4h). These findings clearly show that abJGNs can be eliminated despite the prior acquisition of odor-responsiveness. Notably, the fraction of odor-responding cells among eliminated abJGNs (63.16%) was not significantly different from that measured in surviving abJGNs (56.37%, 146/259 cells, 11 mice) (Fig. 4h, *P* = 0.74, Chi-square test with Yates’ correction), suggesting that acquired odor-responsiveness per se cannot protect abJGNs from being eliminated.

To investigate whether eliminated and surviving abJGNs responded differently to odor stimulation, we analyzed the maximum ΔR/R amplitude and the area under the curve (AUC) of odor-evoked Ca^2+^ transients in eliminated and surviving abJGNs. Then, the median values of eliminated cells were normalized to the median values of corresponding surviving cells recorded from the same mice at the same time points. There was no significant difference between eliminated and surviving cells in terms of the maximum ΔR/R amplitude (Fig. 4i; *P* = 0.49, Wilcoxon signed rank test) and the AUC (Fig. 4j; *P* = 0.23, One sample *t* test; n = 12 odor responding eliminated and 112 odor responding surviving cells, 8 mice). These data demonstrated the lack of correlations between the properties of odor-evoked responses and the fate of abJGNs.

### Distinct endogenous Ca^2+^ signaling in eliminated and surviving abJGNs

Next, we examined whether the Twitch-2B ratio, reflecting the cells’ intracellular free Ca^2+^ concentration ([Ca^2+^]i), or the pattern of endogenous Ca^2+^ signaling can predict the fate of abJGNs. To do so, we analyzed endogenous activity patterns of subsequently eliminated cells at DBD 6 (96-144 hours before death), DBD 4 (48-96 hours before death) and DBD 2 (0-48 hours before cell death) and compared them to median activity patterns of all surviving abJGNs recorded in the same mice at the same time points (Fig. 5). Our previous study described the ubiquitous presence of spontaneous Ca^2+^ transients in the Twitch-2B^+^ abJGNs (Maslyukov et al., 2018). Using tetrodotoxin, a blocker for voltage-gated Na^+^ channels, we established that Twitch-2B^+^ abJGNs can be considered as active (i.e. spiking) when their Twitch-2B ratio is above 2.4 and non-active (i.e. electrically silent) when their Twitch-2B ratio is below 2.0 (Fomin-Thunemann et al., 2020; Maslyukov et al., 2018). In the current study, we used this knowledge to analyze the following parameters: basal Twitch-2B ratio, maximum Twitch-2B ratio, maximum ΔR/R amplitude of spontaneous Ca^2+^ transients, the fraction of time spent in the active state (i.e., Twitch-2B ratio above 2.4), AUC/second (as illustrated in Fig. 5a) and the fraction of time spent in the silent state (i.e., Twitch-2B ratio below 2.0). Many parameters, including the basal (Fig. 5b) and maximum (Fig. 5c) Twitch-2B ratios, the fraction of time spent above 2.4 (Fig. 5e), AUC/second (Fig. 5f) and the fraction of time spent below 2.0 (except DBD 4; Fig. 5g) were elevated in the subsequently eliminated abJGNs (*P* < 0.05 for all comparisons, Wilcoxon signed rank test; DBD 6: n = 14 eliminated and 302 surviving cells from 7 mice, DBD 4: 20 eliminated and 466 surviving cells from 8 mice, DBD 2: 16 eliminated and 371 surviving cells from 8 mice). Only the maximum ΔR/R amplitudes (Fig. 5d) were similar between the subsequently eliminated and surviving abJGNs at all 3 time points.

**Figure 5.**
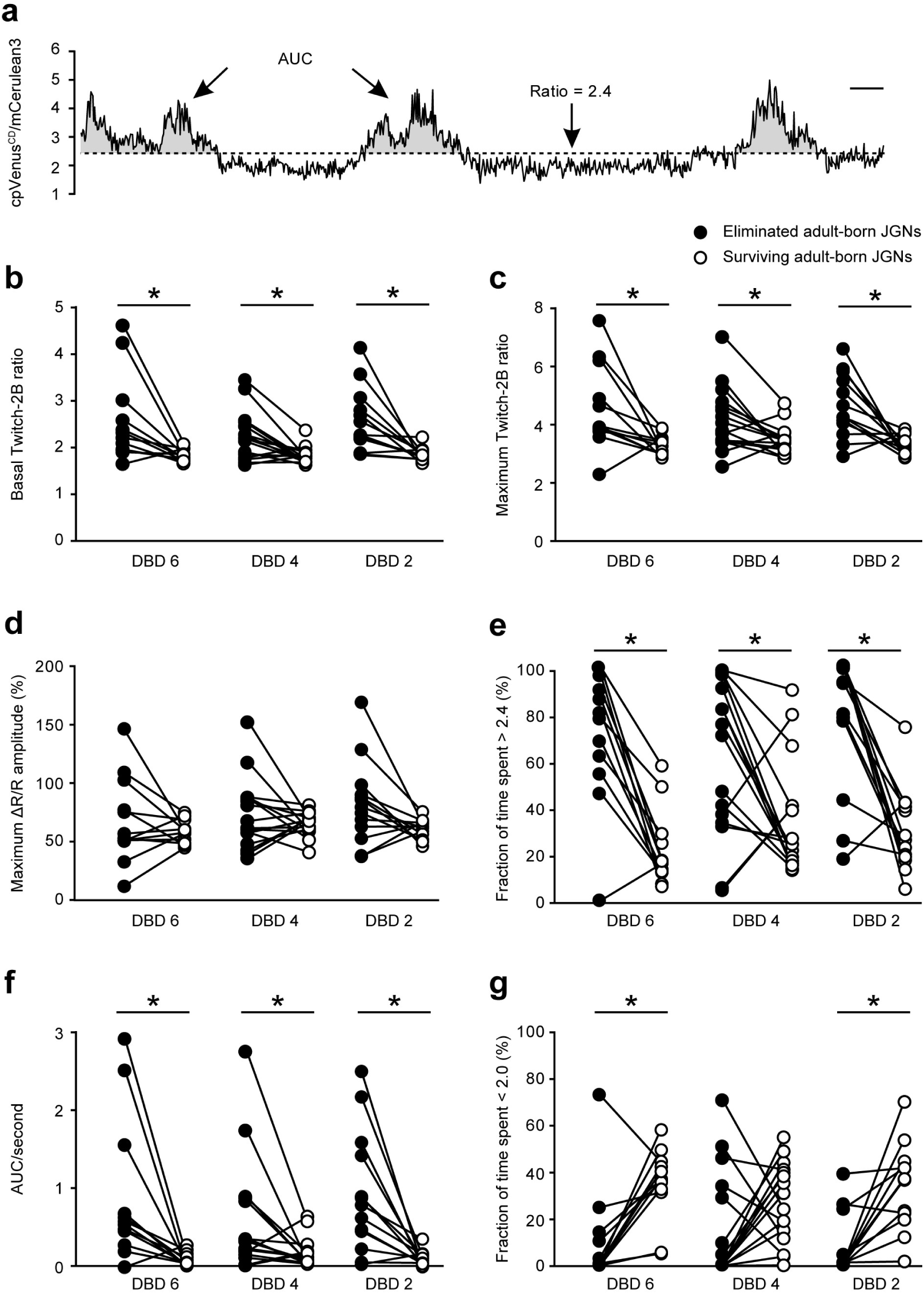
Basal Twitch-2B ratios and endogenous activity patterns differ between eliminated and surviving abJGNs. (**a**) Graph illustrating the endogenous activity of abJGNs and parameters used for its analyses. AUC: area under the curve. Scale bar: 5 s. (**b-g**) Connected dot graphs summarizing the median values of basal Twitch-2B ratio (\**P* < 0.05, Wilcoxon signed rank test) (**b**), maximum Twitch-2B ratio (\**P* < 0.05, Wilcoxon signed rank test) (**c**), maximum ΔR/R amplitude (*P* > 0.05, Paired *t* test) (**d**), fraction of time spent above 2.4 (i.e. in the active state; \**P* < 0.05, Wilcoxon signed rank test) (**e**), the normalized area under the curve (AUC/second; \**P* < 0.05, Wilcoxon signed rank test) (**f**) and fraction of time spent below 2.0 (in the non-active state; \**P* < 0.05, Wilcoxon signed rank test) (**g**) of eliminated and corresponding surviving cells from the same mice recorded at DBD 6, DBD 4 and DBD 2. DBD 6: n = 14 eliminated and 302 surviving cells from 7 mice; DBD 4: 20 eliminated and 466 surviving cells from 8 mice; DBD 2: 16 eliminated and 371 surviving cells from 8 mice. The following supplementary figures are related to Figure 5: **Supplementary Figure 7**. Stability of basal Twitch-2B ratios and endogenous activity patterns in eliminated and surviving abJGNs. **Supplementary Figure 8**. Analysis of [Ca^2+^]_i_ fluctuations in abJGNs.

Next, we investigated the stability of endogenous Ca^2+^ signaling in surviving and subsequently eliminated abJGNs from DBD 6 to DBD 2. To this end, we took DBD6 as a reference point and calculated, for all parameters under study, the differences between values measured at DBD 6 and at subsequent time points (Supplementary Fig. 7). Both for the surviving and eliminated cells, the values measured in individual neurons scattered around zero. The median values for surviving cells were remarkably similar in the three recording sessions (Supplementary Fig. 7; *P* > 0.05 for all comparisons, Friedman test). For eliminated abJGNs, however, we did observe an increase in the basal (Supplementary Fig. 7a) and maximum (Supplementary Fig. 7b) Twitch-2B ratio and the maximum ΔR/R amplitude (Supplementary Fig. 7c) at DBD 2 compared to DBD 4 and DBD 6, probably reflecting the apoptosis-related rise of [Ca^2+^]_i_ (Cellerino et al., 2000; Linden et al., 2005) in some subsequently eliminated cells. However, the detected differences between the surviving and subsequently eliminated abJGNs were not significant (Supplementary Fig. 7; *P* > 0.05 for all comparisons, Wilcoxon signed rank test; n = 8 eliminated cells and 138 surviving cells, 5 mice).

It was suggested that fluctuations of [Ca^2+^]_i_ rather than mere Ca^2+^ influx are needed to drive the phosphorylation of nuclear cAMP-response-element-binding protein (CREB; Li et al., 2016, 2009), a known key mediator of the development and survival of ABCs (Giachino et al., 2005; Jagasia et al., 2009; Li et al., 2016, 2021). To determine whether abJGNs were eliminated because their [Ca^2+^]_i_ was unable to fluctuate, we analyzed the fluctuations of [Ca^2+^]_i_ both in eliminated and surviving abJGNs (see Methods).

At all 3 time points tested (DBD 6, DBD 4, DBD 2), the fractions of cells with fluctuating [Ca^2+^]_i_ did not differ significantly between eliminated and surviving abJGNs (Supplementary Fig. 8; *P* > 0.05 for all comparisons, Chi-square test with Yates’ correction; DBD 6: n = 14 eliminated and 302 surviving cells from 7 mice, DBD 4: 20 eliminated and 466 surviving cells from 8 mice, DBD 2: 16 eliminated and 371 surviving cells from 8 mice), suggesting that the high absolute levels of [Ca^2+^]_i_ rather than the lack of fluctuations were predictive for the subsequent cell death.

Thus, the endogenous activity patterns and the accompanying ongoing changes in [Ca^2+^]_i_ differ significantly between eliminated and surviving abJGNs, with eliminated abJGNs showing substantially increased levels of [Ca^2+^]_i_ over prolonged periods of time.

## Discussion

Using longitudinal intravital imaging, we have characterized the morphological and functional properties of subsequently eliminated and surviving adult-born OB interneurons. According to our conservative estimate, subsequently eliminated cells comprise at least 20% of adult-born JGNs. The elimination is largely accomplished at the end of the pre-integration phase, with no abJGNs being eliminated after DPI 34. Surprisingly, stable surviving abJGNs grew their dendritic trees only during a brief initial time period (DPI 9-13) and spent the majority of the maturation time (from DPI 13 onward) reducing and refining their dendritic trees. Despite the distinct morphological appearance, adult-born PGCs and SACs showed similar patterns of dendritic development. This knowledge could not have been anticipated based on the previous population-based data, describing the net dendritic growth of adult-born JGNs as the major hallmark of their maturation (Livneh et al., 2014, 2009; Mizrahi, 2007). Similar patterns of dendritic morphogenesis were observed in surviving and eliminated abJGNs. Importantly, we found that 63% of the subsequently eliminated abJGNs acquired odor-responsiveness prior to cell death, a rate of odor-responsiveness which was comparable to that of surviving cells. Finally, our data clearly showed that the levels of endogenous Ca^2+^ signaling differ significantly between the surviving and subsequently eliminated abJGNs. However, the death of abJGNs was associated with a protracted increase in the endogenous Ca^2+^ signaling, much in contrast to the previous concept, associating enhanced neuronal activity with increased survival of ABCs (Lin et al., 2010).

The use of a nucleotide analog BrdU as the lineage-tracing tool revealed a significant level of cell death among immature ABCs (∼50% of cells; Petreanu and Alvarez-Buylla, 2002; Winner et al., 2002; Yamaguchi and Mori, 2005). Recently, an elegant *in vivo* study (Platel et al., 2019) has cast doubt on this phenomenon. By using Nestin-CreER^T2^xRosa-RFP mice, the authors injected tamoxifen to induce RFP expression in SVZ stem cells and all their progeny and measured the number of red cells in the OB once a week between week post-injection 1 and 8. Analyzed in this way, only 1.5% of abJGNs and 5.9% of adult-born GCs disappeared over the observation time period. Similar data were obtained in 48 tdTomato^+^ adult-born GCs, labeled by lentiviruses injected into the RMS, prompting the authors to suggest that previously observed cell death is caused by the toxicity of high doses of BrdU (Platel et al., 2019). Although our conservative estimation of the fraction of eliminated abJGNs (∼20%) is also a factor of 2 lower than that found using BrdU, giving room to the phenomenon of BrdU-mediated toxicity, it is still 10 times higher than that reported by Platel et al. The explanation likely lies in the population-based nature of all previous studies. Indeed, when imaging many (20-170) red cells once a week (Platel et al., 2019), it is very difficult to account for vivid cell migration (Liang et al., 2016), the substitution of dying cells by late-coming ABCs (Sawada et al., 2011), etc. Consistently, longitudinal every other day *in vivo* imaging of adult-born GCs revealed that ∼21% of adult-born GCs were eliminated between DPI 24 and DPI 56 (∼0.7%/day decline in total cell number; Sailor et al., 2016). Taken together, these data show that considerable fractions of ABCs are eliminated after the arrival to their destination layers. For abJGNs, the period of cell elimination seems to start at latest at DPI 12, somewhat earlier than assumed previously (Mandairon et al., 2006; Platel et al., 2019), and last till DPI 34 only. This time window closely coincides with the period of lateral migration during the pre-integration phase (Liang et al., 2016).

Migrating immature neurons usually have a highly polarized morphology: a long leading and a short trailing process (Buchsbaum and Cappello, 2019; Ota et al., 2014). This is also true for immature ABCs migrating tangentially in the RMS or radially in the deeper layers of the OB (Doetsch et al., 1997; Kaneko et al., 2017). For adult-born GCs, which stop and settle down at the end of radial migration (Liang et al., 2016), the steady growth of the dendritic tree (Petreanu and Alvarez-Buylla, 2002; Sailor et al., 2016) is a natural next step in the maturation process. The abJGNs, however, after accomplishing the radial migration enter a 1-3 weeks long lateral migration phase, during which they migrate and grow their dendritic trees simultaneously (Kovalchuk et al., 2015; Liang et al., 2016). How these two processes influence each other remained unclear. We show now that abJGNs, which settle down by DPI 13, significantly reduce their dendritic complexity during the subsequent development, removing even primary dendrites (Fig. 2). Although surprising and counterintuitive at first glance, these data suggest that while migrating, abJGNs spread their dendrites around to sense activity patterns of many neighboring glomeruli. Once the parent glomeruli (Kovalchuk et al., 2015) have been found, the cells stop and start pruning unnecessary dendrites, which go beyond their parent glomeruli. This hypothesis can well explain the higher odor-responsiveness but lower odor-selectivity of immature abJGNs, when compared with their mature counterparts (Livneh et al., 2014). A similar concept might also apply to adult-born GCs. These cells also show overshooting of dendritic complexity, albeit to a much lesser extent (Sailor et al., 2016), higher odor-responsiveness and lower odor-selectivity (Wallace et al., 2017) during the maturation process.

Our data show that 63% of abJGNs responded to odor stimuli before their death (Fig. 4h). This number is likely underestimated, as instead of determining for each adult-born JGN the odorant activating its parent glomerulus (Kovalchuk et al., 2015), we simply applied a mixture of odorants broadly activating dorsal glomeruli (Kovalchuk et al., 2015; Livneh et al., 2014) to increase the throughput. Nonetheless, our data demonstrated that the acquisition of odor-responsiveness does not prevent subsequent cell death. The parameters of odor-evoked responsiveness analyzed in this study (maximum response amplitude and AUC), were also not able to differentiate between surviving and subsequently eliminated cells. We cannot, however, exclude the possibility that other properties (e.g., odor selectivity or the stability of odor responses; Livneh et al., 2014; Wallace et al., 2017; Fomin-Thunemann et al., 2020) differ in these two groups of cells. In this study, we took the cells’ odor-responsiveness as a read-out of its functional integration into surrounding neural circuitry. Whether the eliminated odor-responsive cells were indeed synaptically connected, remains unclear. There is, however, a consensus that PSD95-GFP puncta serve as a good proxy for putative synapses (Kelsch et al., 2009, 2008; Livneh et al., 2009). Livneh et al. have shown that dendrites of abJGNs at DPI 12 are already abundantly decorated by such puncta and the number of puncta increases during the maturation process (Livneh et al., 2009). Therefore, our eliminated odor-responsive abJGNs, which were older than DPI 22, likely were synaptically connected (i.e., integrated into the pre-existing circuitry) prior to their elimination.

Interestingly, our data revealed that eliminated abJGNs exhibited elevated endogenous Ca^2+^ signaling over a prolonged time, thus identifying the level of endogenous Ca^2+^ signaling as a potential predictor of the cells’ fate. This is in contrast to the literature data, suggesting that enhanced neuronal activity promotes the survival of immature adult-born GCs (Lin et al., 2010). Although the exact reason for this discrepancy remains to be discovered, it has to be noted that while the NaChBac expression increased the survival of adult-born GCs in the OB (Lin et al., 2010), it failed to do so in adult-born hippocampal GCs (Sim et al., 2013), suggesting that different types of ABCs might possess different survival strategies (Pfisterer and Khodosevich, 2017). In addition, we observed the enhanced endogenous activity in subsequently eliminated ABCs after DPI 18 while in the previous study (Lin et al., 2010) the NaChBac expression started at ABCs’ birth. Thus, timing may also affect the outcome.

Having discovered that the subsequent death of abJGNs can be predicted by the level of their endogenous Ca^2+^ signaling, we hypothesized that this is due to the loss of fluctuations in [Ca^2+^]_i_, needed for optimal activation of the CREB-mediated survival program (Li et al., 2016, 2009). Surprisingly, the vast majority of subsequently eliminated abJGNs maintained fluctuations in [Ca^2+^]_i_ even at DBD 2, right before undergoing apoptosis (Supplementary Fig. 8). Although we cannot exclude the possibility that some subtle fluctuation features, which are difficult to extract because of the inhomogeneity of the activity patterns in individual Twitch-2B^+^ cells, do differ between the surviving and the eliminated cells, our current data suggest that the death of abJGNs is triggered by the sustained increase in the overall level of [Ca^2+^]_i_. In this respect, mitochondria might be an important player. A sustained increase in [Ca^2+^]_i_ causes a sustained increase in the Ca^2+^ content of the endoplasmic reticulum (ER) (Garaschuk et al., 1997) and might promote a direct (via the mitochondrial Ca^2+^ uniporter or voltage-dependent anion channel) or an ER-mediated (via mitochondria-associated ER membranes) sustained Ca^2+^ accumulation in the mitochondria (Celsi et al., 2009; Rizzuto et al., 2012). The latter can trigger oxidative stress and the release of proapoptotic factors (e.g. cytochrome c, apoptotic inducing factor, procaspase 9, endonuclease G) into the cytosol (Calvo-Rodriguez et al., 2020; Giorgi et al., 2012).

Taken together, our study identifies endogenous Ca^2+^ signaling as a key player influencing the fate of abJGNs. The latter can integrate both the cell-intrinsic firing and the influence of the steady-state odor environment reaching either a pro- or an anti-apoptotic level. Because some odorants increase and others decrease the levels of endogenous activity in JGNs (Homma et al., 2013), this model reconciles the seemingly opposing findings showing that odor enrichment either decreases (Khodosevich et al., 2013) or increases (Forest et al., 2019; Rey et al., 2012; Rochefort et al., 2002) the survival of abJGNs.

## Materials and methods

### Animals

All experiments were designed and conducted in accordance with biometrical planning and institutional animal welfare guidelines and were approved by the government of Baden-Württemberg, Germany. Three- to four-month-old C57BL/6 mice of either sex were used in this study. Animals were kept in pathogen-free conditions at 22 °C and 60% air humidity, under 12 hours light/dark cycle with *ad libitum* access to food and water. Female mice were maintained in groups of 3-5 mice, male mice were maintained individually. All experimental procedures followed the Directive 2010/63/EU of the European Parliament and the Council of the European Union.

### Cranial window implantation

A cranial window above the mouse OB was implanted as described previously (Kovalchuk et al., 2015; Liang et al., 2016; Maslyukov et al., 2018). Briefly, mice were anesthetized by an intraperitoneal (i.p.) injection of ketamine (Fagron, Barsbuettel, Germany) and xylazine (Sigma-Aldrich, MO, USA; 80/4 μg/g body weight (BW)). The depth of anesthesia was estimated using a toe pinch reflex and an additional dose of ketamine/xylazine (40/2 μg/g BW) was injected when necessary. Mice were head-fixed in a stereotaxic setup (Stoelting, IL, USA) and a circular groove (Ø 3 mm) was made above the OB using a high-speed dental drill (ultimate 500, NSK). After careful removal of the bone with fine tweezers (leaving the dura intact), a glass coverslip (Ø 3 mm, Warner Instruments, CT, USA) was gently positioned above the OB. Cyanoacrylate glue was applied to the gap between the coverslip and the skull to stabilize the coverslip, which was further strengthened by blue light-cured dental cement (Ivoclar Vivadent AG, Liechtenstein). After surgery, analgesic carprofen (5 μg/g BW, Pfizer, Berlin, Germany) was injected subcutaneously for 3 consecutive days and antibiotic enrofloxacin (Baytril, 1:100 v/v, Bayer, Leverkusen, Germany) was applied in drinking water for 10 days.

### Stereotaxic viral injection into the RMS and holder implantation

Approximately 4 weeks after window implantation, stereotaxic virus injection was performed as described previously (Kovalchuk et al., 2015; Liang et al., 2016; Maslyukov et al., 2018). A glass capillary (tip diameter ∼30-40 μm) filled with 1.0-1.5 μl of virus-containing solution (composition see below) was navigated to one of the two injection sites at the following coordinates: AP 3.0mm, ML ± 0.82 mm, DV -3.0 ± 0.05 mm from pial surface. The viral solution contained either a mixture of retroviruses encoding mCherry, Venus and Cerulean (1:1:1 ratio in terms of infectious titer; Gomez-Nicola et al., 2014; Liang et al., 2016) or lentivirus encoding the Ca^2+^ indicator Twitch-2B (Kovalchuk et al., 2015; Maslyukov et al., 2018; Thestrup et al., 2014). After injection, a custom-made titanium holder was fixed to the skull with blue light-cured dental cement. After surgery, analgesic carprofen (5 μg/g BW, Pfizer, Berlin, Germany) was injected subcutaneously for 3 consecutive days.

### Longitudinal *in vivo* two-photon imaging

A training period, allowing the mice to accustom to the experimenter and the setup, started at DPI 5 (Maslyukov et al., 2018). The mice were gently handled inside home cages and then transferred to the microscope stage for 30 minutes of free exploration. At DPI 6-11, after gentle handing and free exploration the mice were head-fixed within the setup for 5 (at DPI 6) to 30-40 (at DPI 11) minutes. Longitudinal *in vivo* two-photon imaging in awake mice began at DPI 12.

Image acquisition was performed with a customized two-photon microscope based on an Olympus Fluoview 1000 system (Olympus, Tokyo, Japan), coupled to a mode-locked Ti:Sapphire laser (Mai Tai Deep See, Spectra Physics, CA, USA), as described previously (Homma et al., 2013). The emitted fluorescence light was collected through a 20x water immersion objective (NA 1.0, Carl Zeiss, Jena, Germany). Three-dimensional (3D) stacks (512x512 pixels, Kalman filter 2, step 3 µm) were acquired daily at DPI 12-35 and DPI 44-45 to document the locations of RGB-positive (RGB^+^) cells in awake mice. We used 800 nm wavelength to excite Cerulean and mCherry and a 570-nm dichroic mirror (570 DM) to split the emission light (the short-pass channel was used for Cerulean and the long-pass channel for mCherry) as well as 970 nm wavelength to excite Venus (emitted fluorescence was collected via the short-pass channel of 570 DM). Because each labeled cell had its color code, this procedure allowed a precise tracking of individual migrating cells (Gomez-Nicola et al., 2014; Liang et al., 2016). Sparse labeling, blood vessel patterns, tiny irregularities of the dura and the distance between the stable RGB^+^ cell were used as landmarks helping to identify the same cells in longitudinal imaging sessions (Gonçalves et al., 2016a).

To image the dendritic morphology of RGB^+^ abJGNs, mice were head-fixed under isoflurane anesthesia (2% for induction, 0.8-1.2% for maintenance) to prevent any animal movements. Both eyes were protected with eye ointments (Bepanthen, Bayer, Germany). Body temperature was maintained at 36-37 °C and breath rate was kept at 100-140 breaths per minute (BPM). We chose the excitation wavelength (900 or 970 nm), optimally exciting the expressed fluorescent proteins. The 3D stacks (512x512 pixels, Kalman filter 3, step 2 µm) were recorded with zoom factors, which were as high as possible and allowed to include the entire dendritic tree into the FOV.

### *In vivo* two-photon Ca^2+^ imaging

Starting from DPI 11, mice were trained 7 consecutive days for awake imaging, as described above. At DPI 18-44, 3D stacks (512x512 pixels, Kalman filter 2, step 3 µm) were acquired at 930 nm excitation wavelength every second day to document the locations of Twitch-2B^+^ cells. Endogenous Ca^2+^ signals were imaged (sampling rate: 6.67 Hz) in awake mice at 890 nm excitation wavelength. Fluorescence signals from mCerulean and cpVenus^CD^ were split at 515-nm and filtered with a 475/64 nm band-pass filter for mCerulean and a 500 nm long-pass filter for cpVenus^CD^ channels, respectively (Semrock, Rochester, United States).

### Odor application

Odor-evoked responses were measured as described in (Homma et al., 2013). In brief, mice were anesthetized with an i.p. injection of midazolam (5 mg/kg BW), medetomidine (0.5 mg/kg BW) and fentanyl (0.05 mg/kg BW). Body temperature was maintained at 36-37 °C and breathing rate was 130-170 BPM. A custom-made flow dilution olfactometer was used to deliver a mixture consisting of the equal parts of either 3 (isoamyl acetate, 2-hexanone and ethyl tiglate) or 7 (ethyl-acetate, butanal, pentanal, ethyl tiglate, propanal, methyl-propionate and ethyl-butyrate) odorants, causing broad activation of dorsal OB glomeruli (Kovalchuk et al., 2015; Livneh et al., 2014). The mixture of odorants (1.7% of saturated vapor) was presented as a 4-second-long pulse in front of the mouse’s snout.

### Data analyses

For the analysis of dendritic morphology, neuronal soma and dendrites were manually traced with Simple Neurite Tracer plugin of Image J (https://imagej.nih.gov/ij). Cells with incomplete dendritic arborizations (e.g. located at the edge of the field of view) were excluded from the analysis. Total dendritic branch length (TDBL), as well as numbers of dendrites, primary dendrites, branch points and endings, were quantified with Neuromantic (Myatt et al., 2012) and Analyze Skeleton plugin of Image J. We considered dendrites emerging from the soma as primary dendrites, points of dendritic bifurcation as branch points (Ledderose et al., 2014; Lee et al., 2013), dendrites between the dendrite tip and a branch point (Gonçalves et al., 2016a) as endings (see also Fig. 2a). Sholl analyses were performed using Simple Neurite Tracer plugin in Image J. Pruned and added dendrites between the 2 consecutive time points were analyzed with the computer-aided 4D structural plasticity analysis (4DSPA) software, according to the established procedure (Gonçalves et al., 2016a; Lee et al., 2013). Results were further processed in MATLAB (The MathWorks, Inc., Massachusetts, USA).

For the analysis of SBR and SNR, for each cell 5 dendritic endings were chosen in Fiji. For each dendritic ending, SBR was calculated as the ratio of its mean fluorescent intensity to the mean fluorescent intensity of the corresponding background (darkest area near the cell). SNR was defined as the ratio of ending’s mean fluorescent intensity to the standard deviation of the fluorescent intensity of the background. SBR (or SNR) of a cell equaled the median of the SBRs (or SNRs) of its 5 dendritic endings. For analyses of the endogenous activity, a somatic region of interest (ROI) and a background ROI (the darkest area near the target cell) of the comparable size were drawn in Image J. The readouts of mCerulean3 and cpVenus^CD^ fluorescence intensity were imported into MATLAB and filtered with a 1st order low-pass digital Butterworth filter with a cutoff frequency of 0.6 Hz. The Twitch-2B ratio was calculated using the following formula:

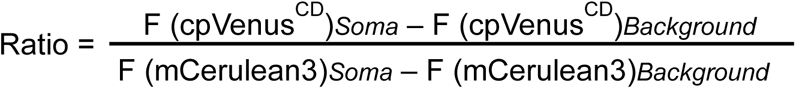

The following parameters were analyzed: basal Twitch-2B ratio, maximum Twitch-2B ratio, maximum ΔR/R amplitude, fraction of time spent above the ratio of 2.4, area under the curve (AUC), fraction of time spent below the ratio of 2.0. Basal and maximum Twitch-2B ratios were calculated as follows: the filtered traces were processed in MATLAB using a sliding average algorithm with a window size of 5 seconds to determine the basal Twitch-2B ratio (minimum average value) and 1.5 seconds to determine the maximum Twitch-2B ratio (maximum average value). The amplitude (ΔR/R) was calculated as (R-R_0_)/R_0_, where R_0_ is the basal Twitch-2B ratio. In fluctuation analysis, we included the Twitch-2B ratio traces of each eliminated and all (more than 10) corresponding surviving cells, recorded at the same time point from the same mice. In some mice, there were several cells, eliminated at different time points. In this case, we analyzed the Twitch-2B ratio traces of the same surviving abJGNs taken at time points, corresponding to DBD 6, 4 and 2 of the respective eliminated cell. Fluctuations of the Twitch-2B ratio, reflecting fluctuations in the JGN’s [Ca^2+^]_i_, were detected using the mid-reference level crossing approach (IEEE Instrumentation and Measurement Society, 2011). The degree of fluctuation has been quantified using the number of crossing points passing through the mid-reference level. The Gaussian mixture model (McLachlan and Peel, 2000) has been used to detect the naturally existing clusters in the counted crossing points. The correct number of clusters has been estimated using Bayesian Information Criteria (Wit et al., 2012). Assigned labels have been used to identify the cells belonging to each cluster among eliminated and corresponding surviving abJGNs. Obtained fractions were compared statistically using the Chi-square test with Yates’ correction. For the analyses of odor-evoked responses, the amplitude was calculated using a custom-written routine in Igor Pro (WaveMetrics, Portland, USA). All ΔR/R traces were filtered using a binomial filter (time window 0.3 s) and the filtered traces were subtracted from the original ΔR/R traces to obtain the trace of background noise. Cells were defined as odor-responding if the maximum ΔR/R amplitude of their odor-evoked signals was 3 times larger than the standard deviation of the corresponding background noise.

### Statistical analyses

Unless indicated, all data are shown as median ± interquartile range (IQR). Statistical analyses were performed using GraphPad Prism 7 (GraphPad Software, CA, USA). Shapiro-Wilk test was used to check the normality of data distribution in individual datasets. For analyses of a single dataset, One sample *t* test or Wilcoxon signed rank test was applied. For two unpaired datasets, either Unpaired *t* test or Mann-Whitney test were used, depending on the normality of the datasets. For two paired datasets, Paired *t* test or Wilcoxon signed rank test were applied. For 3 or more paired datasets, Friedman test followed by Dunn’s multiple comparisons test or One-way repeated measures ANOVA with or without Greenhause-Geisser Sphericity Correction followed by Holm-Sidak post hoc test was applied. Unless indicated, all statistical tests were two-tailed. Differences were considered significant if *P* < 0.05.

## Acknowledgments

We thank J. Tiago Goncalves and Yu-Tai Ching for their help with 4DSPA analysis, E. Zirdum, A. Weible and K. Schoentag for excellent technical assistance, K. Li and K. Figarella for viral production, N. Asavapanumas for his help with endogenous activity analysis, A. Gohl, J. Mueck, M. Knecht and S. Wang for help with morphological analysis, S. Jessberger, Y. Liang and K. Li for critical comments on the manuscript. This work was funded by the DFG grant (GA 654/14-1) to O.G.; X.S. was partially supported by the DAAD Graduate School Scholarship Program (Personal Reference Number: 91586010).

## Author contributions

O.G. initiated and conceived the study. X.S. and Y.K. performed experiments. X.S. analyzed the data. N.M. contributed to data analyses. X.S. and O.G. wrote the manuscript. All authors have approved the final version of the manuscript.

---

**Supplementary Figure 1.**
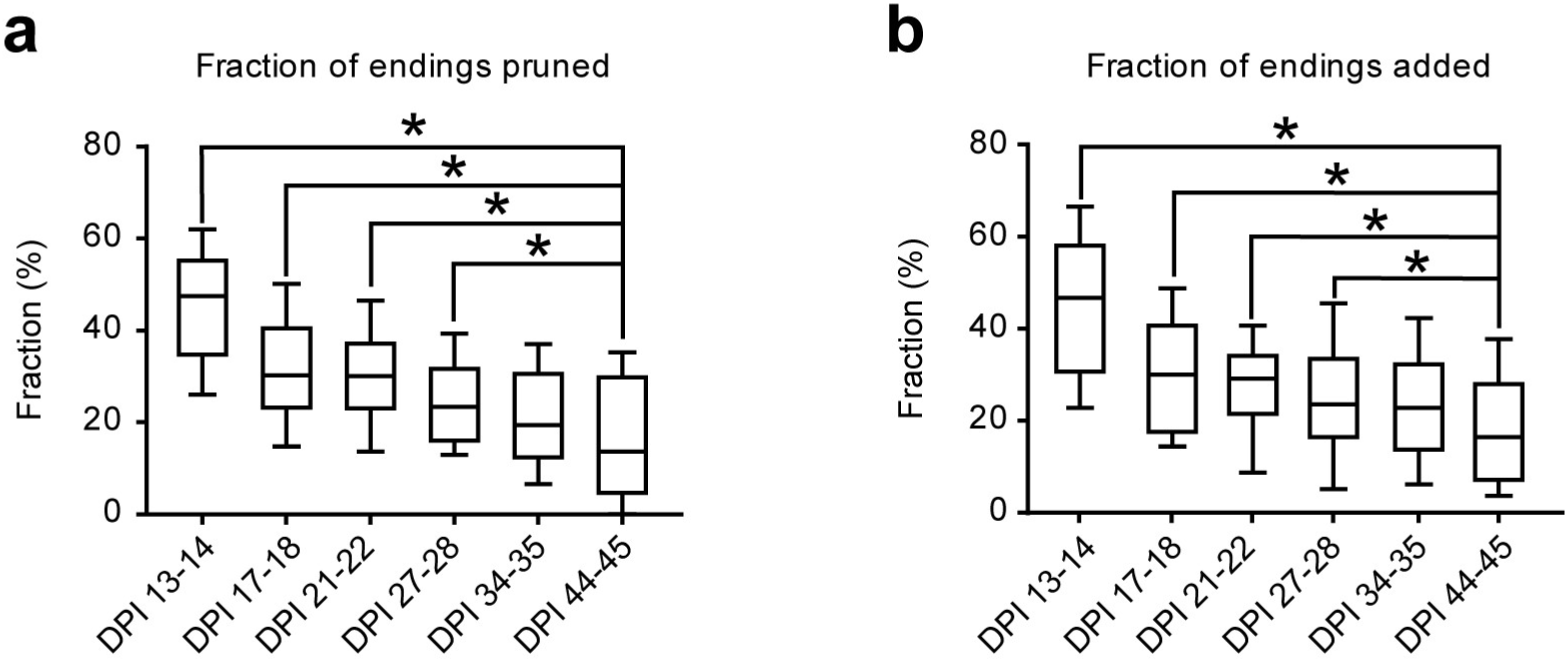
4D structural plasticity analysis of dendritic remodeling in stable surviving abJGNs. (**a,b**) Box plots showing the fractions of pruned (**a**) and added (**b**) dendritic endings in stable surviving abJGNs (\**P* < 0.05, One-way repeated measures ANOVA followed by Holm-Sidak’s multiple comparisons test; n = 34 cells, 8 mice).

**Supplementary Figure 2.**
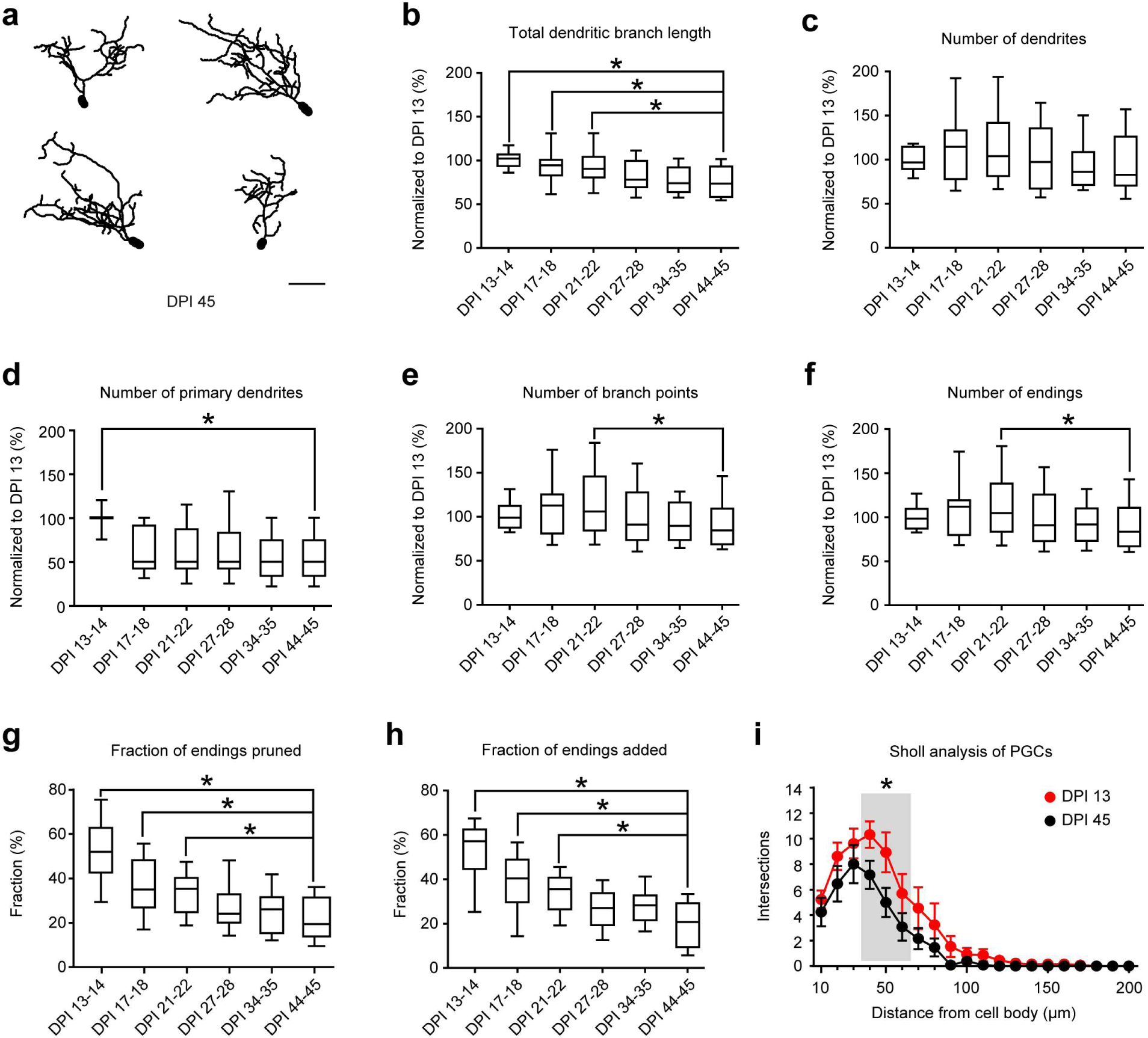
Development and plasticity of the dendritic tree in stable surviving adult-born PGCs. (**a**) Representative reconstructions of 4 adult-born PGCs at DPI 45. Scale bar: 25 µm. (**b-f**) Box plots summarizing the normalized total dendritic branch length (**b**; *\*P* < 0.05, One-way repeated measures ANOVA followed by Holm-Sidak’s multiple comparisons test) as well as the numbers of dendrites (**c**; *P* > 0.05, One-way repeated measures ANOVA followed by Holm-Sidak’s multiple comparisons test), primary dendrites (**d**; *\*P* < 0.05, Friedman test followed by Dunn’s multiple comparisons test), branch points (**e**; *\*P* < 0.05, One-way repeated measures ANOVA followed by Holm-Sidak’s multiple comparisons test) and endings (**f**; *\*P* < 0.05, One-way repeated measures ANOVA followed by Holm-Sidak’s multiple comparisons test) in stable surviving adult-born PGCs. For each cell, the data were normalized to the respective values measured at DPI 13 (n = 13 PGCs, 7 mice). (**g, h**) 4D structural plasticity analysis showing the fractions of pruned (**g**) and added (**h**) endings in stable surviving adult-born PGCs (*\*P* < 0.05, One-way repeated measures ANOVA followed by Holm-Sidak’s multiple comparisons test; n = 13 PGCs, 7 mice). (**i**) Sholl analyses of adult-born PGCs at DPI 13 (red) and DPI 45 (black). Grey zone marks the region, where the numbers of intersections showed significant differences between the two age groups (*\*P* < 0.05, Wilcoxon signed rank test, n= 13 PGCs, 7 mice). Error bars: SEM.

**Supplementary Figure 3.**
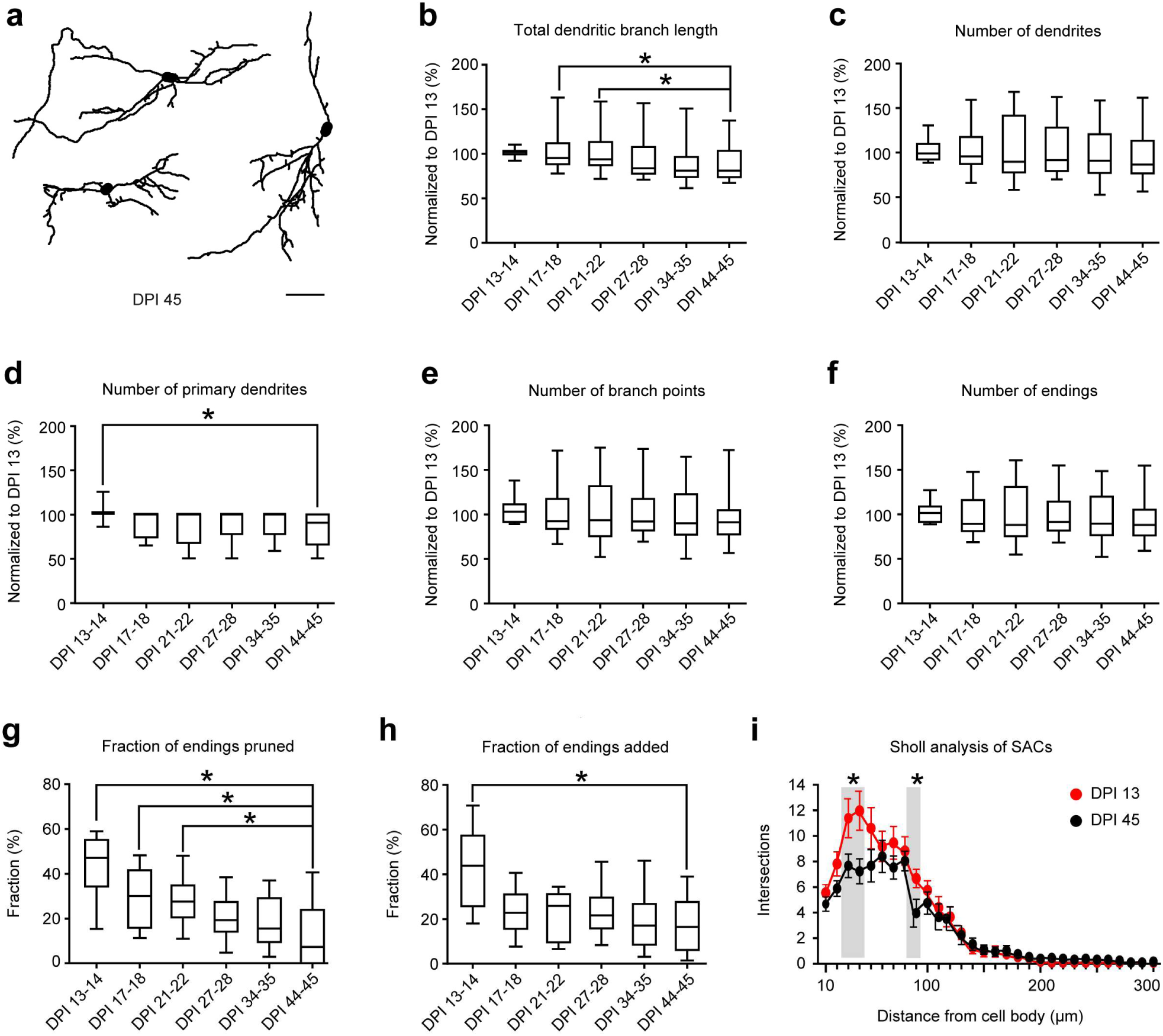
Development and plasticity of the dendritic tree in stable surviving adult-born SACs. (**a**) Representative reconstructions of 3 adult-born SACs at DPI 45. Scale bar: 50 µm. (**b-f**) Box plots illustrating the normalized total dendritic branch length (**b**; *\*P* < 0.05, One-way repeated measures followed by Holm-Sidak’s multiple comparisons test) and the numbers of dendrites (**c**; *P* > 0.05, Friedman test followed by Dunn’s multiple comparisons test), primary dendrites (**d**; *\*P* < 0.05, Friedman test followed by Dunn’s multiple comparisons test), branch points (**e**; *P* > 0.05, Friedman test followed by Dunn’s multiple comparisons test), and endings (**f**; *P* > 0.05, One-way repeated measures ANOVA followed by Holm-Sidak’s multiple comparisons test) in stable surviving adult-born SACs. For each cell, the data were normalized to the respective values measured at DPI 13 (n = 14 SACs, 5 mice). (**g, h**) 4D structural plasticity analysis showing the fractions of pruned (**g**) and added (**h**) dendritic endings at DPI 13-45 (*\*P* < 0.05, Friedman test followed by Dunn’s multiple comparisons test; n = 14 SACs, 5 mice). (**i**) Sholl analyses of SACs at DPI 13 (red) and DPI 45 (black). Grey zones mark the regions, where the numbers of intersections showed significant differences between the two age groups (*\*P* < 0.05, Wilcoxon signed rank test; n = 14 SACs, 5 mice). Error bars: SEM.

**Supplementary Figure 4.**
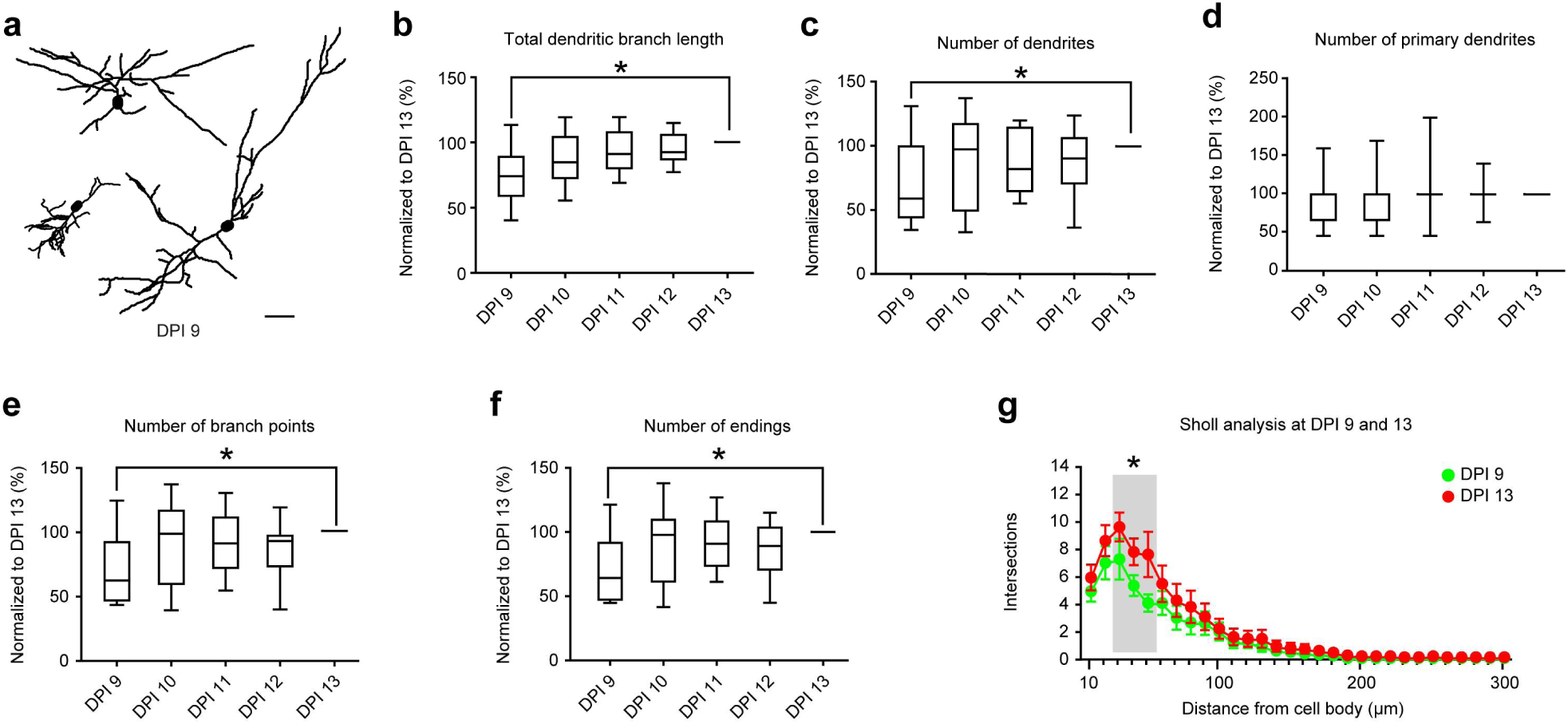
Initial period of rapid dendritic growth in surviving abJGNs at DPI 9-13. (**a**) Representative reconstructions of 3 abJGNs at DPI 9. Scale bar: 25 µm. (**b-f**) Box plots summarizing the normalized total dendritic branch length (**b**) as well as the numbers of dendrites (c), primary dendrites (**d**), branch points (**e**) and endings (**f**) in surviving abJGNs at DPI 9-13. For each cell, the data were normalized to the respective values measured at DPI 13 (*\*P* < 0.05 for all comparisons, Friedman test followed by Dunn’s multiple comparisons test; n = 15 JGNs, 4 mice). (**g**) Sholl analyses, showing the number of dendritic intersections with Sholl spheres at DPI 9 (green) and DPI 13 (red). Grey zone marks the region, where the numbers of intersections showed significant differences between the two age groups (*\*P* < 0.05, Wilcoxon signed rank test; n = 15 JGNs, 4 mice). Error bars: SEM.

**Supplementary Figure 5.**
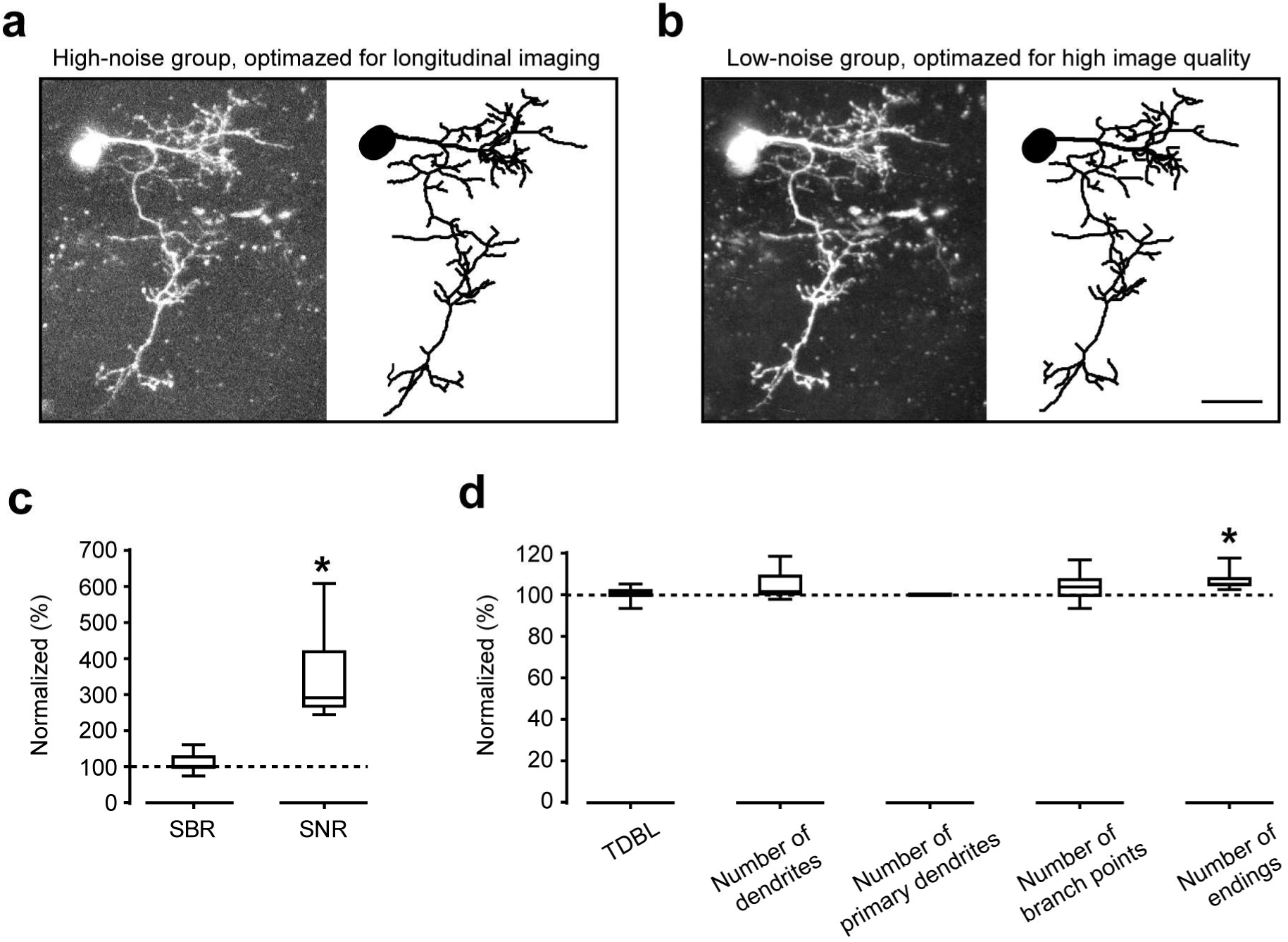
Impact of the improved image quality on the measurements of dendritic complexity of abJGNs. (**a, b**) Representative MIP images (14-52 µm, step 2 µm) and 3D reconstructions of the same abJGN imaged under high (**a**) and low (**b**) noise conditions. Scale bar: 20 µm. **(c**) Box plots showing the normalized signal-to-background (SBR, left) and signal-to-noise (SNR, right) ratios (*\*P* < 0.001, Wilcoxon signed rank test, n = 12). For each cell, the SBR (or SNR) in the low-noise group was normalized to the respective SBR (or SNR) in the high-noise group. (**d**) Box plots summarizing the normalized TDBL as well as the numbers of dendrites, primary dendrites, branch points and endings (*\*P* < 0.001, Wilcoxon signed rank test, n = 12). For each cell, the data in the low-noise group were normalized to the respective values in the high-noise group.

**Supplementary Figure 6.**
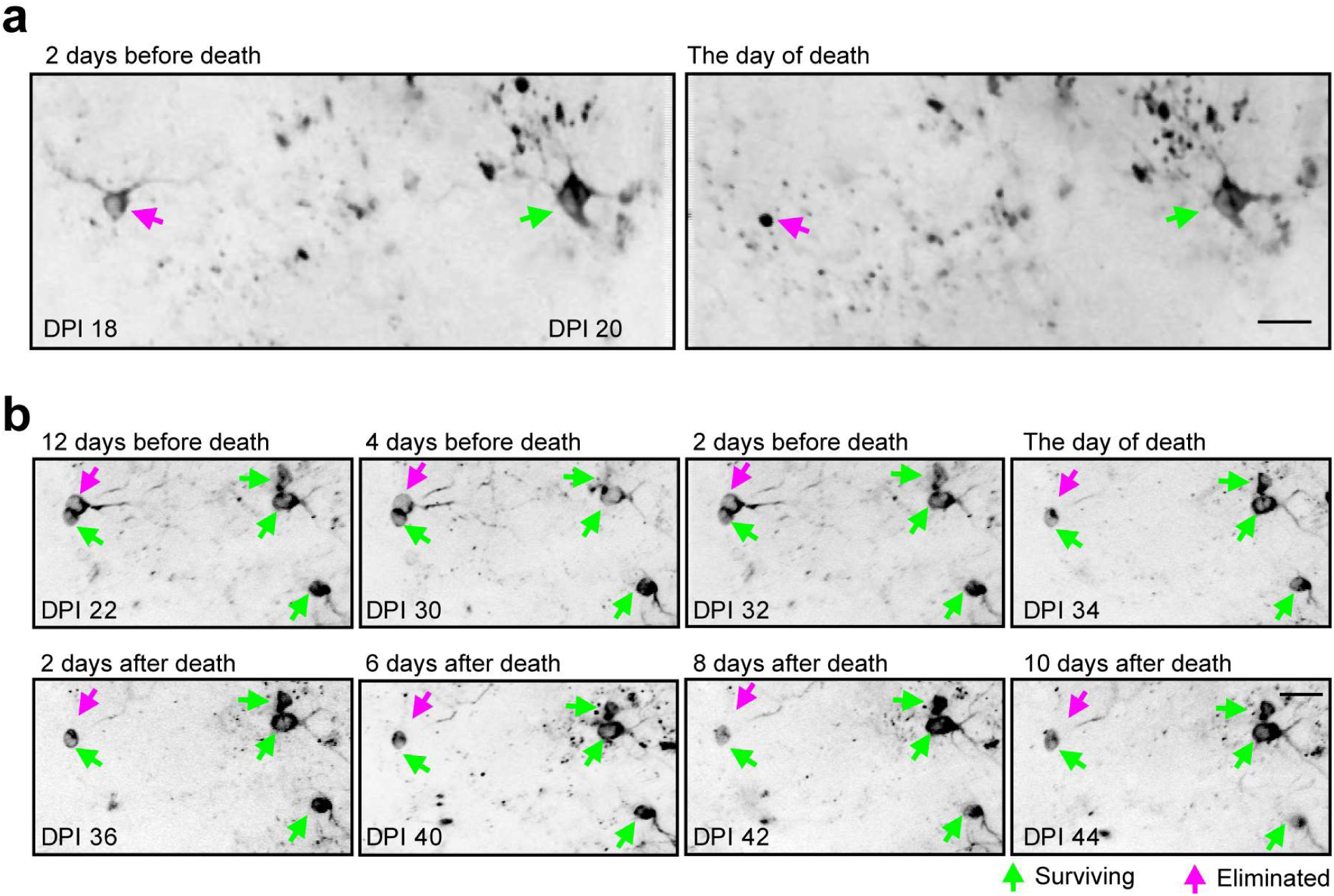
Visualization of the death of Twitch-2B^+^ abJGNs. (a) Images showing 1 eliminated (magenta arrow) and 1 surviving (green arrow) Twitch-2B^+^ abJGNs in the same FOV at DPI 18 (left) and DPI 20 (right). Here and in (b) each image is an average of 830 consecutive frames. Scale bar: 20 μm. (b) Images showing 1 eliminated (magenta arrow) and 4 surviving (green arrows) cells in the same FOV at 8 different time points. DPI 34 was defined as the day of death. Scale bar: 20 μm.

**Supplementary Figure 7.**
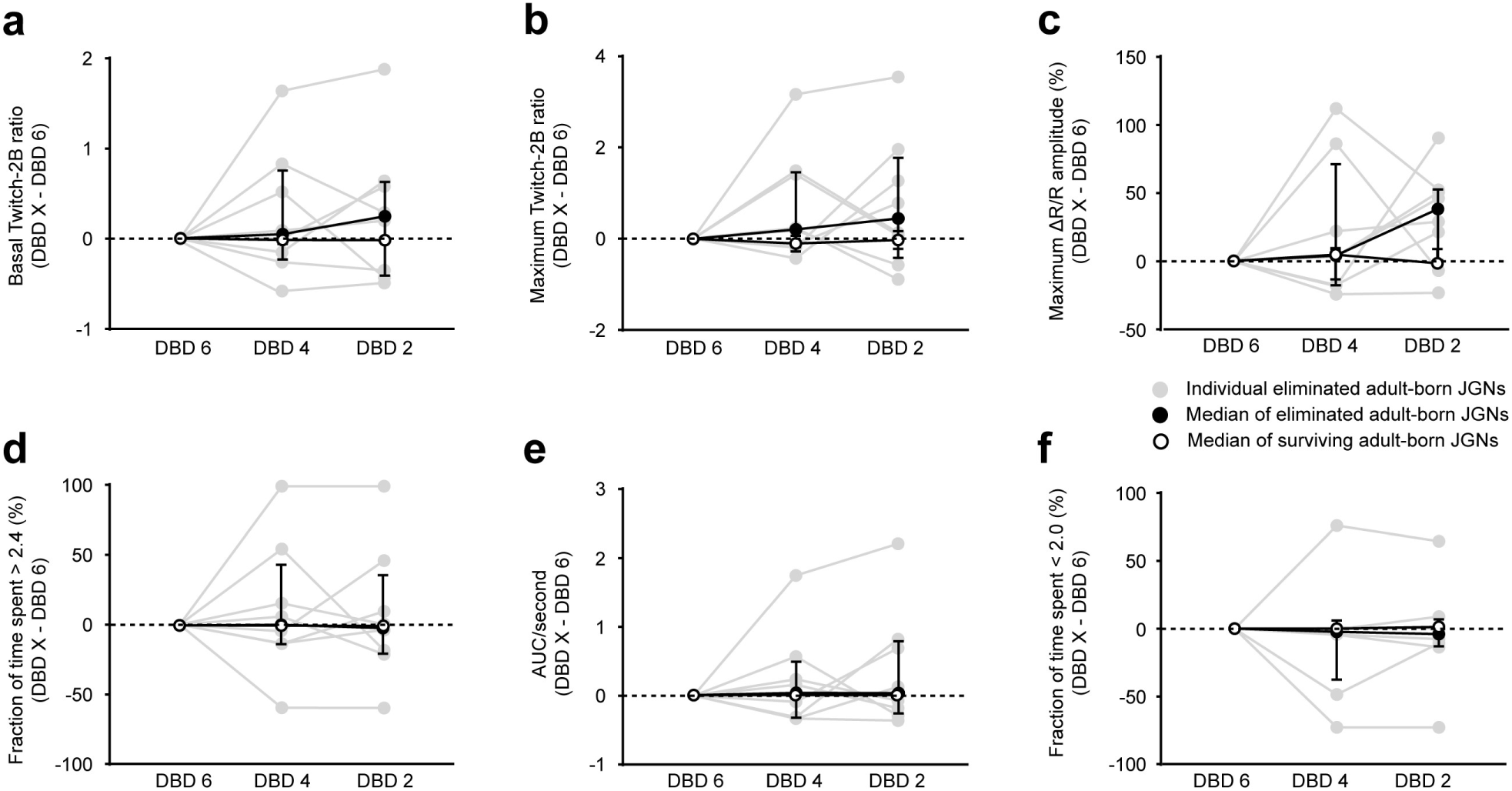
Stability of basal Twitch-2B ratios and endogenous activity patterns in eliminated and surviving abJGNs. (**a-f**) Graphs, showing the differences between DBD 6 and DBD 6, DBD 4 or DBD 2 for the following parameters: basal Twitch-2B ratio (**a**), maximum Twitch-2B ratio (**b**), maximum ΔR/R amplitude (**c**), fraction of time spent above 2.4 (**d**), normalized AUC (AUC/second) (**e**), fraction of time spent below 2.0 (**f**). Different symbols correspond to different cell groups as indicated in (**c**). *P* > 0.05 for all comparisons, Friedman test for comparison of the same groups at different time points, Wilcoxon signed rank test for comparison of different groups at the same time points; n = 8 eliminated cells and 138 surviving cells, 5 mice.

**Supplementary Figure 8.**
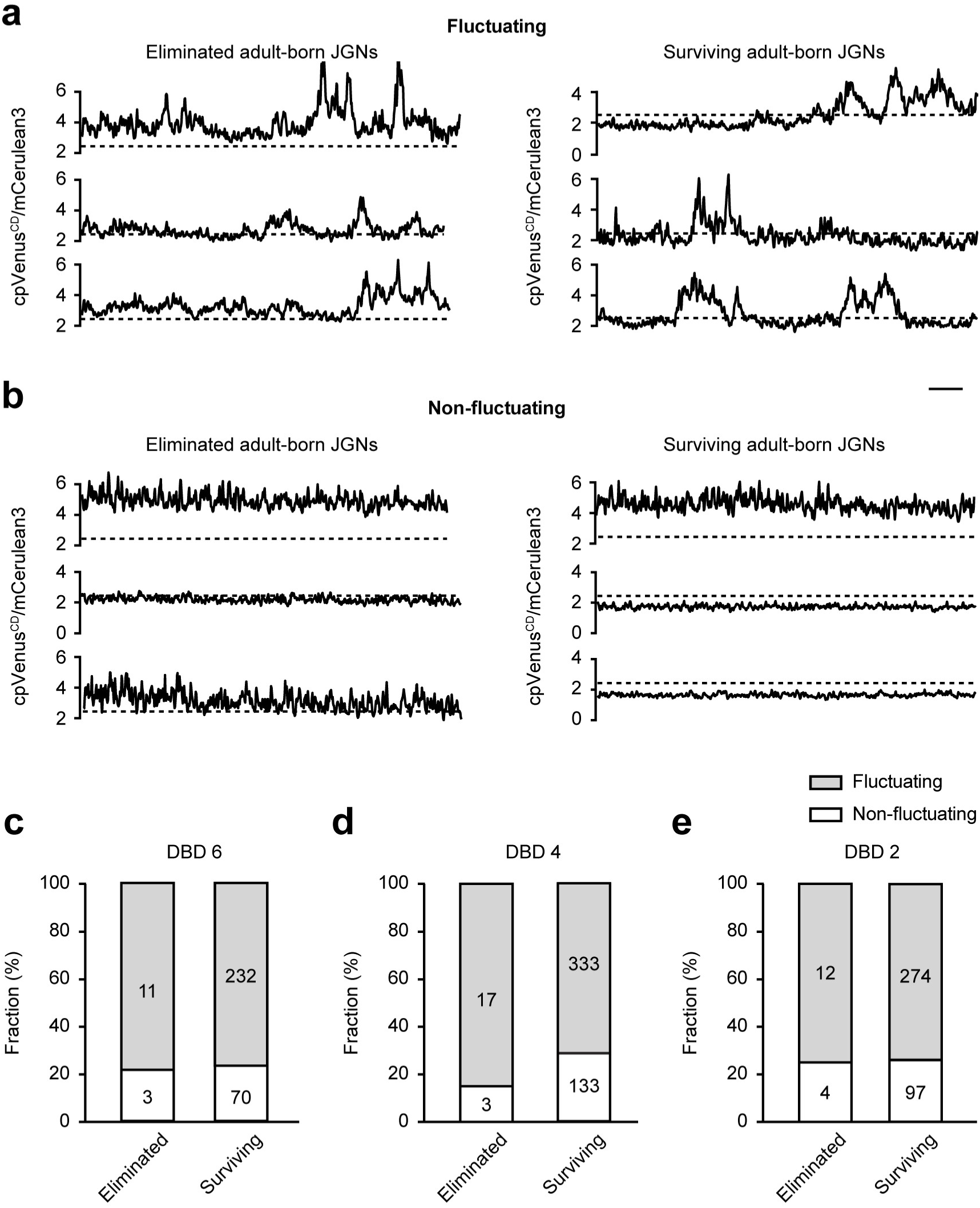
Analysis of [Ca^2+^]_i_ fluctuations in abJGNs. (**a, b**) Representative traces, obtained from abJGNs with fluctuating (**a**) and non-fluctuating (**b**) Twitch-2B ratios. Scale bar: 10 s. (c-e) Bar graphs summarizing the fractions of cells with fluctuating and non-fluctuating Twitch-2B ratios among the subsequently eliminated and surviving abJGNs at DBD 6 (**c**), DBD 4 (**d**) and DBD 2 (**e**). *P* > 0.05 for all comparisons, Chi-square test with Yates’ correction.

